# Data-adaptive three-dimensional deconvolution and evaluation for volumetric fluorescence microscopy

**DOI:** 10.64898/2026.06.29.735443

**Authors:** Yiwei Hou, Yunzhe Fu, Wenyi Wang, Ruijie Cao, Xuantao Su, Donghyun Kim, Meiqi Li, Peng Xi

## Abstract

Optical fluorescence microscopy enables visualization of biological structures and dynamics. However, the intrinsic diffraction limit, especially axially, and depth-related scattering noise compromise the image resolution and fidelity. Computational 3D deconvolution is a promising approach for mitigating these issues, yet its execution is hindered by inaccurate and cumbersome theoretical modeling or experimental measurement of 3D point spread function (PSF), as well as ineffective 3D noise regularization. Furthermore, in the 3D super-resolution regime, there remains a lack of standardized tools for evaluating 3D super-resolution fidelity. Here, we present the 3D adaptive deconvolution and evaluation (3D-ADE) toolkit, which comprises 3D-Ada deconvolution with physics-oriented automatic 3D-PSF calibration, and 3D-SQUIRREL for 3D super-resolution quality assessment. It effectively resolves noise instability, eliminates the need for 3D-PSF calibration, and reliably assesses the fidelity of 3D resolution extension via deconvolution, physical, and deep-learning-based methods. Accessible via multiple software platforms, 3D-ADE enhances the versatility of 3D deconvolution and fills the gap in 3D super-resolution evaluation tools, and thereby advances volumetric fluorescence imaging applications.

## Introduction

Optical fluorescence microscopy^1^ is a cornerstone technique for visualizing biological structures and dynamics. However, a fundamental physical constraint limits its performance in three-dimensional (3D) imaging: the microscope objective typically can only collect back-propagating fluorescence signals. This inherently leads to lower axial than lateral resolution, making isotropic 3D visualization challenging. This anisotropy persists even in state-of-the-art 3D imaging techniques, including confocal^2-4^, multi-photon^5, 6^, and light-field microscopy^7, 8^, as well as super-resolution methods like stimulated emission depletion (STED)^9^ and structured illumination microscopy (SIM)^10-13^, compromising image fidelity.

To recover fine details from the blurred 3D raw data, computational deconvolution is a powerful approach. Yet, executing 3D-deconvolution faces profound challenges, primarily centered on accurately and conveniently determining the 3D point-spread-function (PSF). Compared to 2D modeling, theoretical modeling of 3D-PSF requires a larger set of imaging parameters and is more sensitive to distortions from case-dependent optical aberrations and sample-induced refractive index mismatch^14^. These practical imperfections are difficult to incorporate into a static theoretical PSF model based on manually recorded imaging parameters. Experimentally measuring the 3D-PSF is cumbersome^15^, requires high expertise, is susceptible to noise^16^, and is uncertain on inter-sample transferring^14^. This “PSF dilemma” renders most existing 3D deconvolution methods unstable in practice and poses a major obstacle to pursuing isotropic resolution through deconvolution. Furthermore, despite noise regularization substantially boosting the performance of 2D-deconvolution^17-19^, these models are less effective in 3D due to anisotropic sampling, resolution, and more complex noise distributions. Current 3D-deconvolution approaches are primarily rudimentary inverse iterations techniques that may lack noise robustness^20-22^.

Furthermore, there is a notable absence of standardized analytical tools for evaluating artifacts in 3D-deconvolution, physical^3, 12, 23-26^, and the emerging deep-learning-based 3D super-resolution^27-29^, hindering objective assessment of result fidelity.

To address these problems, we introduce the 3D adaptive deconvolution and evaluation (3D-ADE) toolkit, offering an image-driven automatic 3D-PSF determination, noise-robust 3D deconvolution, and 3D super-resolution artifact evaluation. It aims to enhance the versatility and accessibility of 3D-deconvolution in state-of-the-art fluorescence microscopy techniques, while filling the gap in 3D super-resolution error evaluation.

## Results

### Image-based automatic calibration of the 3D-PSF

In our computation pipeline, robust and versatile noise suppression for 3D data is needed for accurate image analysis. Consequently, we develop an adaptive noise suppression model (**Note S1**) leveraging 3D-Framelet^17, 30^ and Stein’s unbiased estimate of risk (SURE)^31^ mathematical tools. Given a decomposed 3D-Framelet coefficient subset *c*^*i*^, its SURE-criterion thresholding value is calculated as follows:

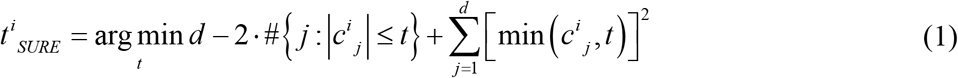

where # denotes the number of elements that satisfy the condition within the braces, *t*^*i*^_*SURE*_ is the thresholding value decided by the SURE criterion, and *d* is the total element number in a coefficient subset. Then the weight of each coefficient subset is calculated as *w*^*i*^ based on its respective *t*^*i*^_*SURE*_ value. Then, a weighted thresholding for noise suppression is performed as follows:

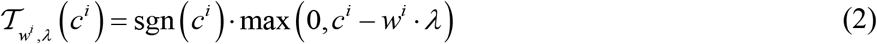

where *λ* is the global thresholding value. We term this noise suppression scheme the Ada noise model. This design accounts for sample and anisotropic 3D sampling characteristics, reflected in the power of the coefficients in different 3D-Framelet sub-bands. It is well adapted for accurate noise suppression in complex 3D structures compared with the commonly adopted uniform thresholding (**Fig. S1**). Ada obtains evidently higher quality metrics over static spatial-variation-based methods, and more importantly, preserves the high-frequency information that is crucial for further analysis (**Figs. S2, 3**).

Leveraging the proposed Ada noise model and advanced Fourier-ring-correlation (FRC) based decorrelation^32^ resolution estimation, we develop an automatic image-driven 3D-PSF estimation method (**Fig. 1a, Note S2, Table S1**). Our core idea is that with a known PSF shape but unknown parameters (for example, the *σ* parameter in a Gaussian shape), once the resolution of the image (i.e., the blur level) can be calculated, the PSF can be obtained by deducing the PSF parameter using the resolution value. Especially, very recently, some advanced resolution estimation methods, such as decorrelation^32^ analysis, have been proposed. We argue that these resolution estimation methods also serve as an effective module in achieving automatic 3D-PSF estimation.

**Fig. 1.**
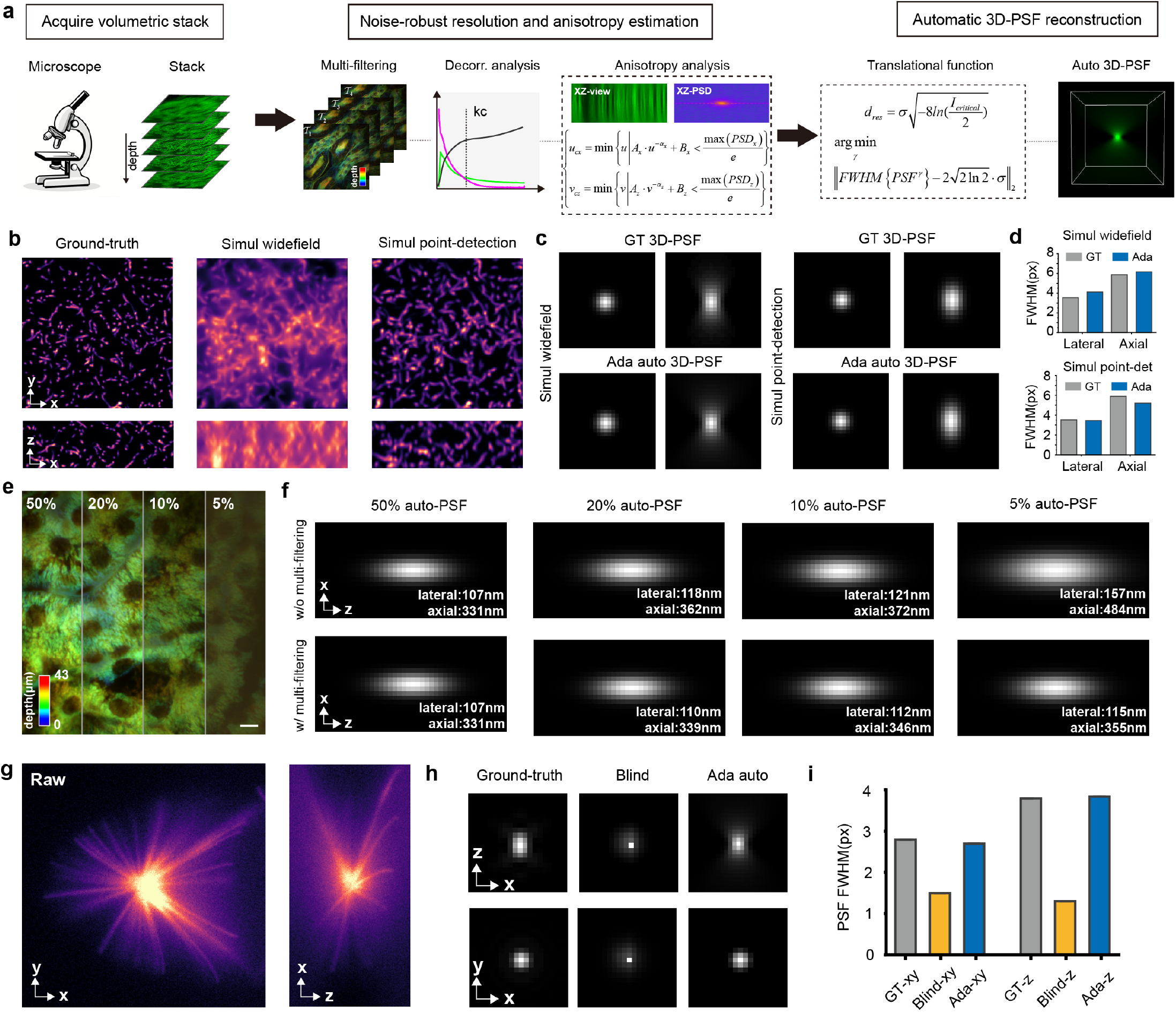
The 3D-Ada deconvolution method. **a**. The flowchart of the Ada automatic 3D-PSF calibration process. **b**. An open-source simulation pattern^29^, and its degenerated forms using a widefield and point-detection 3D-PSF, respectively. **c**. The ground-truth (GT) 3D-PSF and the estimated results of our Ada method. **d**. The FWHM values of the 3D-PSF. **e**. Raw captured SD-confocal images with gradient illumination power. **f**. The Ada estimated 3D-PSF results with and without the devised multi-filtering strategy. The FWHM values are shown on the bottom right. **g**. An open-source microtubule-like pattern. **h**. The GT 3D-PSF, and the estimation results via blind deconvolution and our Ada method. **i**. The FWHM values of the 3D-PSF. Scale bar: 10 *μ*m (e).

Mathematically, we propose transition functions that bridge the connection between resolution value and key shape parameters for 3D-PSF estimation. Given an estimated resolution value *d*_*res*_ obtained via decorrelation analysis, and the critical midpoint intensity value to distinguish two points *I*_*critical*_, we deduce that the *σ* parameter of a Gaussian PSF has the following relationship with the two values (**Note S2**):

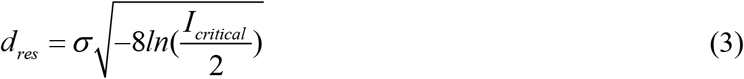

For other PSF shapes, their parameters can be determined by the following criteria that match the full-width-of-half-maxima (FWHM) with a standard Gaussian PSF:

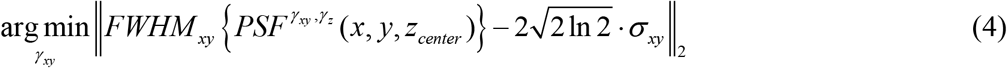

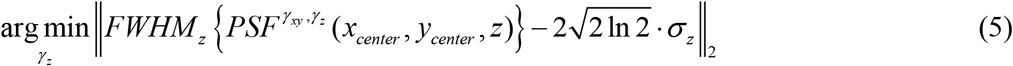

where *γ*_*xy*_ and *γ*_*z*_ denote the parameter that determines the lateral and axial FWHM of the 3D-PSF.

For ease of use and enlightenment from previous studies, we uniformly employ a Gaussian-Lorentzian-shape 3D-PSF for a widefield detection system^33^, and a Gaussian-shape 3D-PSF for a point-detection system^34^ (**Note S2**).

We examine the performance of this pipeline in simulated data in both widefield and point-detection 3D-PSF, all showing very satisfactory estimations that are close to the ground-truth 3D-PSFs (**Fig. 1b, c**). Noise may disturb the decorrelation-based resolution determination and influence the 3D-PSF estimation pipeline. By developing a multi-filtering strategy based on the proposed Ada noise model (**Note S2**), we finely address this issue, ensuring feasibility under practical noise conditions (**Fig. 1e-f, Fig. S4**).

We evaluated the proposed physically grounded 3D-PSF calibration method using the EPFL open-source dataset^22^ with well-calibrated PSFs. Our method provides reliable estimations, with key FWHM parameters close to the calibrated ones in all instances in the dataset (**Fig. 1g-i, Fig. S5**). While the optimization-driven blind deconvolution method^35^ tend to show large deviations of 3D-PSF in FWHM values and shapes.

### 3D-Ada deconvolution

With a given 3D-PSF, we devise an energy-based de-PSF solver incorporating 3D-Ada noise regularization (**Fig. 2a**). It ensures the fidelity of the deconvolution results, contrasting with 3D-Richardson-Lucy (RL) based methods (**Note S3, Fig. S6**). After the noise characteristics analysis, the optimization problem can be formulated as follows:

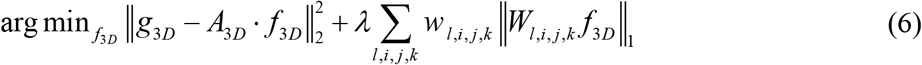

where *g*_3D_ and *f*_*3D*_ denotes the recorded and deconvolved volumetric images respectively, *A*_*3D*_ is the matrix form of the 3D-PSF, *W* denotes the 3D-framelet transform operation, and *w* denotes the weight factor. We extend the original Fast-Iterative-Shrinkage-Threshold-Algorithm (FISTA)^36^ to 3D to solve the optimization problem. The algorithm, combined with the automatic 3D-PSF calibration module developed in the last section, is termed 3D-Ada deconvolution (**Table S2**).

**Fig. 2.**
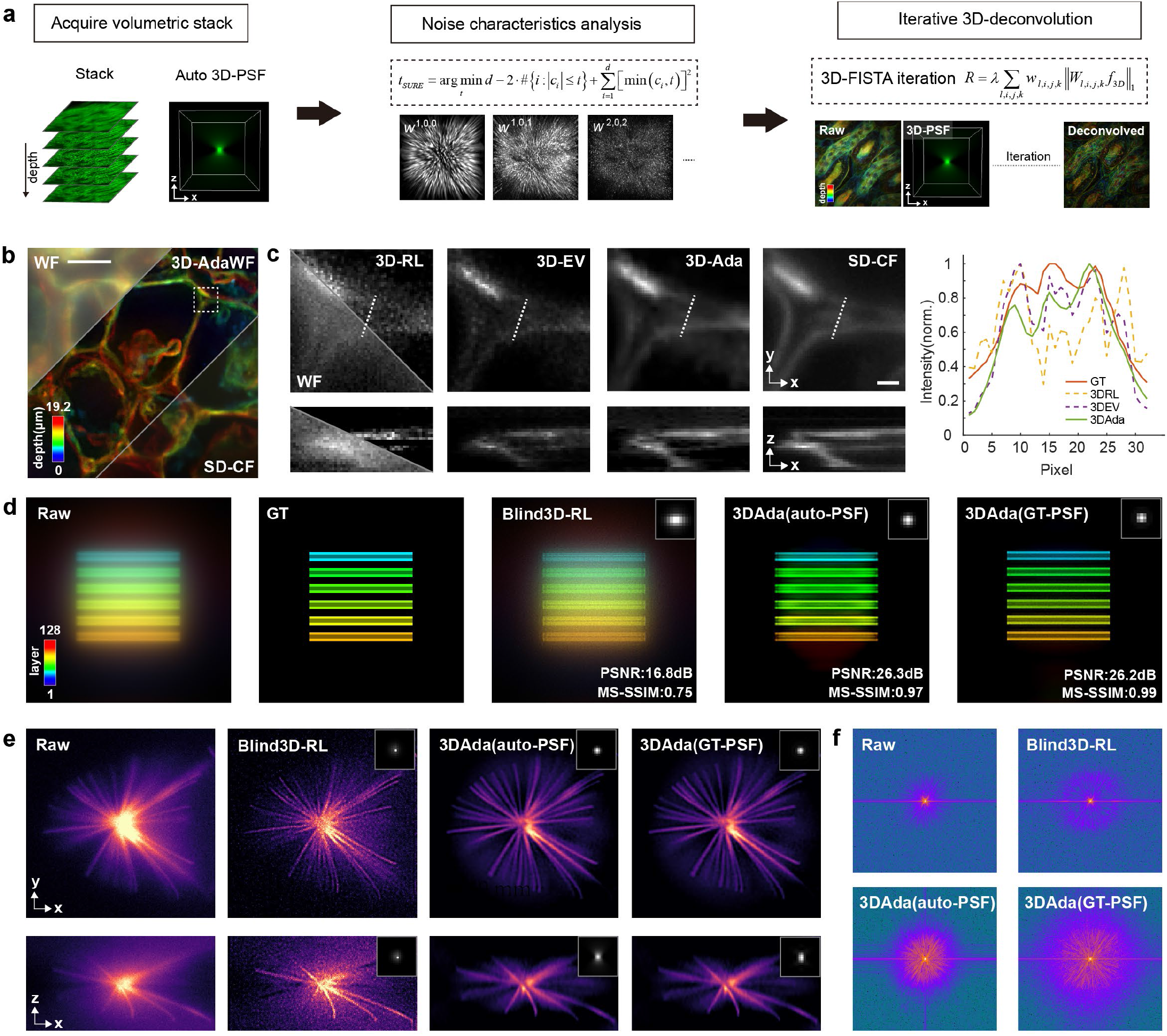
The 3D-Ada deconvolution method. **a.**The flowchart of executing 3D-Ada deconvolution. **b**. Matched widefield and SD-Confocal image of a *Nerlum* stem, and the 3D-Ada deconvolution result. **c**. Magnified white boxed region in b, and the deconvolution results of different algorithms. The xz-slices are made from the centerline. **d**. The “bars” sample in the open-source EPFL dataset, the results produced from blind 3D-RL deconvolution, 3D-Ada deconvolution with automated 3D-PSF and with ground-truth 3D-PSF. **e**. The “microtubule-challenge” sample in the open-source EPFL dataset, the results produced from blind 3D-RL deconvolution, 3D-Ada deconvolution with automated 3D-PSF and with ground-truth 3D-PSF. **f**. Fourier spectra of the average intensity projection in e. Scale bars: 10 *μ*m (b), 1 *μ*m (c).

We first compare the performance of 3D-Ada with state-of-the-art methods, which are 3D-RL and its improved, regularized variant 3D-EV^33^ in cases with known 3D-PSF. In a simulation volume pattern with known ground-truth, 3D-Ada provides finer restoration of the 3D-pattern that can be verified in the ground-truth 3D pattern (**Fig. S7**). While the 3D-RL and 3D-EV methods lack effective signal-noise distinction and over-sharpen the volume, causing noticeable artifacts. We also examine the performance of the algorithms when applied to widefield data using a spinning-disk (SD) confocal data as a reference. Our 3D-Ada deconvolution method demonstrates superior performance over 3D-RL and 3D-EV, effectively suppressing out-of-focus background while recovering true subtle structural details (**Fig. 2b, c**). Besides the known-3D-PSF conditions, we also compare the performance of 3D-Ada with blind deconvolution. The blind deconvolution typically provides very limited recovery performance due to a much-deviated 3D-PSF (**Fig. 2d-f**). 3D-Ada provides satisfactory recovery results with pure automatic 3D-PSF calibration, showing less deviation from the deconvolution results obtained using the ground-truth 3D-PSF.

Together with the automatic 3D-PSF calibration, 3D-Ada enables robust deconvolution for mainstream fluorescence microscopies, especially in situations where PSF calculation or measurement is challenging (**Table S3**). We first show the application of 3D-Ada in deconvolving volumetric light-field microscopy data. The light-field microscopy introduces modulation to enable fast recording of a volume structure. However, its modulation process requires a complicated recalibration of the 3D-PSF that is different from a standard widefield setup. 3D-Ada provides adaptive 3D-PSF in this case (**Fig. 3a**) and provides a fine deconvolution enhancement, while the blind 3D-RL method has a limited recovery effect (**Fig. S8**). We also demonstrate the usage of 3D-Ada in a two-photon microscope when imaging a thick *Pinus radiata* sample^5^ (**Fig. 3b**). 3D-Ada frees the need to model the 3D-PSF in the complex two-photon process, enabling fine augmentation of the raw volume that cannot be achieved through the blind deconvolution method (**Fig. S8**). Besides the two cases that require complicated 3D-PSF calibrations, 3D-Ada can also be conveniently applied to the commonly used SD-confocal setup, enhancing its optical sectioning capability and improving its lateral and axial resolution. With the assistance of 3D-Ada, subtle structures such as the dense actin^15^ (**Fig. 3c**) and the microtubules during the developmental process of *Drosophila* eggs (**Fig. 3d**) can be resolved. 3D-Ada can also be conveniently applied to a super-resolution system with its adaptive 3D-PSF calibration and noise-robustness, like instantSIM (**Fig. 3e**) and 2D-SIM (**Fig. 3f**). The application of 3D-Ada can further improve the 3D-resolution and sectioning capability of the two setups.

**Fig. 3.**
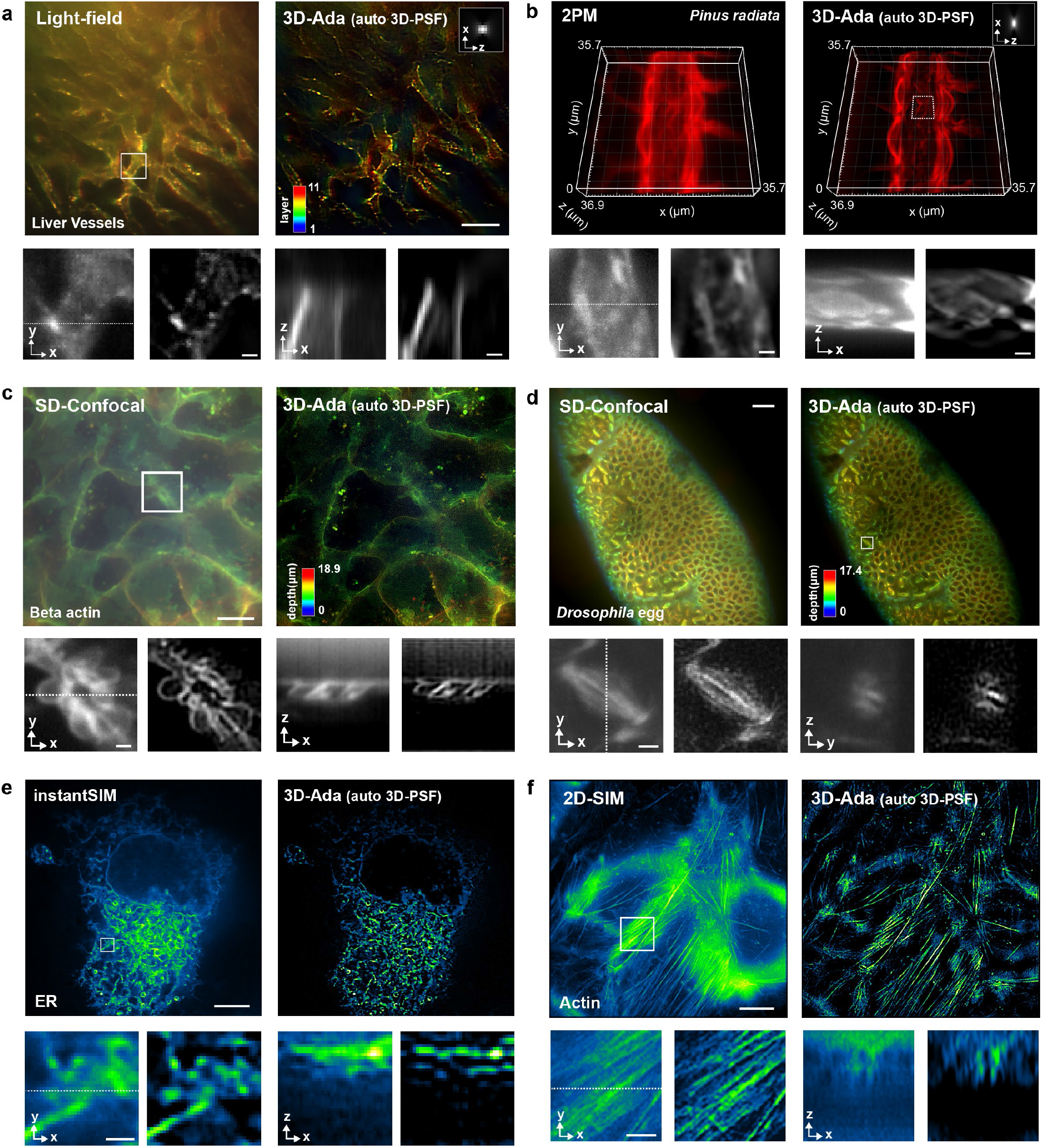
Application of the 3D-Ada deconvolution method. **a**. An open-source liver vessel image captured by a light-field microscope, and its 3D-Ada deconvolution result. **b**. An open-source *Pinus radiata* image captured by a two-photon microscope (2PM), and its 3D-Ada deconvolution result. **c**. An open-source beta actin SD-confocal image, and its 3D-Ada deconvolution result. **d**. A Drosophila egg SD-confocal image, and its 3D-Ada deconvolution result. **e**. An open-source ER image stack captured by an instantSIM microscope, and its 3D-Ada deconvolution result. **f**. An actin stack captured by 2D-SIM, and its 3D-Ada deconvolution result. The automatically estimated 3D-PSFs in a and d are attached in the graphs. Scale bars: 100 *μ*m (a), 10 *μ*m (a subgraph, c, d), 1 *μ*m (e, c subgraph, e subgraph), 2 *μ*m (d subgraph), 5 *μ*m (e, f), 0.5 *μ*m (f subgraph).

### 3D-super-resolution quality evaluation tool

The SQUIRREL (super-resolution quantitative image rating and reporting of error locations) algorithm^37^ quantifies practical super-resolution uncertainties by using a low-resolution reference, which is especially valuable in practical applications without ground-truth. However, it is limited to planar imaging and does not account for axial resolution enhancement. A critical challenge in transferring this method to 3D is the instability of the heuristic parameter search due to its inherent randomness. Here, we solved this issue and developed 3D-SQUIRREL (**Fig. 4a, Note S4**). The full optimization problem is formulated as follows:

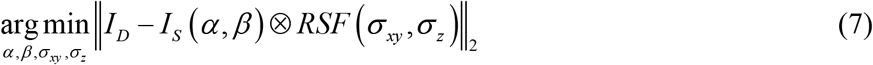

where *I*_*D*_ is a resolution-limited volumetric stack, *I*_*S*_ is a resolution-enhanced volumetric stack, RSF is the 3D resolution-scaled-function parameterized by *σ*_*xy*_ and *σ*_*z*_, *α* and *β* are linear scaling coefficients.

**Fig. 4.**
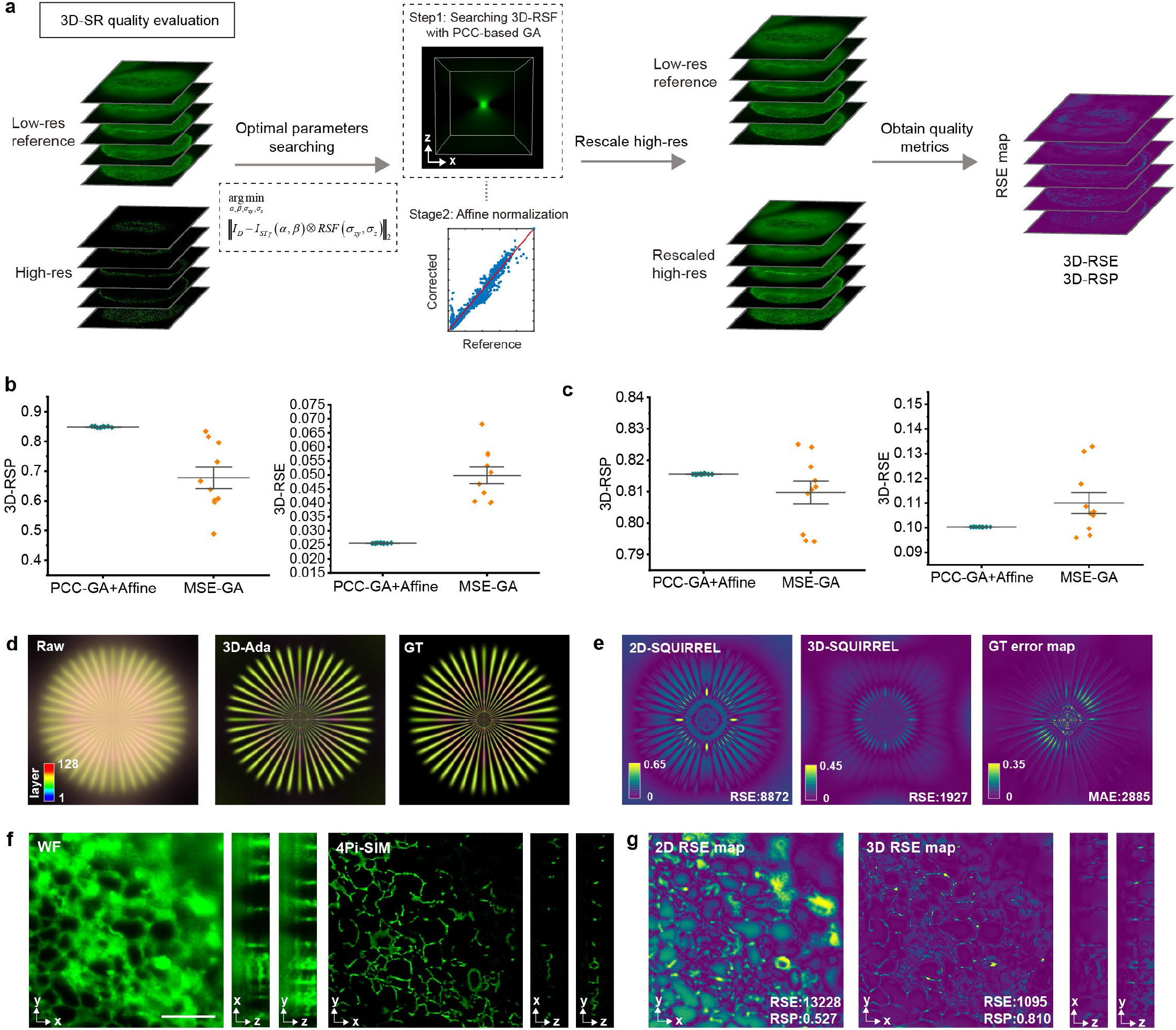
Development of 3D-SQUIRREL and its evaluation. **a**. The computing procedure of the 3D-SQUIRREL algorithm. Robustness test of randomness in the proposed GA searching method for the 3D-SQUIRREL algorithm on **b**. Deconvolved light-field data in Fig. 3a. **c**. Physical 3D-SR results in f. **d**. Application of 3D-SQUIRREL on a representative deconvolved simulation image stack. **e**. The 2D-RSE maps obtained via the original 2D-SQUIREL algorithm, the 3D-RSE maps obtained via the 3D-SQUIRREL algorithm developed here, and the GT error map. **f**. The open-source widefield and 4Pi-SIM image of ER. **g**. the error map given by 2D-SQUIRREL and 3D-SQUIRREL. Scale bar: 5 *μ*m (f).

Our approach decouples the search process into two stages: first, a Pearson correlation coefficient (PCC)-driven parameter search for the 3D-RSF using a genetic algorithm^38^ (GA), followed by a second step of affine normalization to minimize the mean-square-error (MSE). The proposed PCC-GA is essential in a 3D scenario since the rudimentary one-stage MSE-based searching strategy adopted in 2D-SQUIRREL turns out to be rather unstable and often fails to find a near-optimal parameter set (**Fig. 4b, c**). For practical guidance in assessing 3D resolution-enhanced stacks, it can be recommended to accept results with a 3D resolution-scaled-PCC (3D-RSP) greater than 0.8, indicating a strong linear correlation in statistics with the raw recordings^39^.

We first inspect the performance of 2D-SQUIRREL and 3D-SQUIRREL in assessing the volumetric 3D-resolution enhanced data using a simulation pattern with known ground-truth (**Fig. 4d**), and a low-high resolution volume pair of the ER tubule (**Fig. 4f**), which has a structure that is known well and relatively simple. In the simulation, 3D-pattern 3D-SQUIRREL accurately reflects the axial resolution enhancement information and coincides with the true GT-error maps (**Fig. 4d, e**). In contrast, the previous 2D-SQUIRREL may misinterpret axial resolution enhancement as an error. This phenomenon can also be observed in assessing the ER volumetric stack (**Fig. 4f, g**), where the 2D-SQUIRREL would wrongly take the axial resolution enhancement from 4Pi-SIM^26^ as errors. 3D-SQUIRREL can consider the 3D causal and give a confidential 3D super-resolution quality map. This evidence highlights the necessity of using the 3D model for 3D stack evaluation.

### Assessing 3D-super-resolution quality with 3D-SQUIRREL

3D-SQUIRREL serves as an effective tool to assess the quality and fidelity of 3D-resolution enhancement techniques, including 3D-deconvolution, physical 3D-super-resolution microscopy, and 3D-deep-learning-based super-resolution. 3D-SQUIRREL helps guarantee reasonable 3D-deconvolution results that respect the information from raw recordings (**Fig. S9, Table S4**). The 3D-RSP serves as a criterion to judge whether excessive regularization priors or noise amplification are introduced in the deconvolution results.

Additionally, 3D-SQUIRREL can quantify the quality of physical 3D-SIM super-resolution microscopy^12, 23, 40^, revealing the necessary photon-dosage (**Fig. 5a-c**) for reliable 3D-super-resolution. It can also help aid the optimization of the newly developed 3D-super-resolution optical setup (**Fig. S10**). Deep-learning-based methods have emerged as an alternative approach for 3D super-resolution, while assessing their credibility in real applications without ground-truth is still underexplored. 3D-SQUIRREL can judge the reliability of deep-learning-based 3D super-resolution results, offering a solution to this issue. The quality map provided by 3D-SQUIRREL exhibits high similarity with the ground-truth map (**Fig. 5d, e)**, showing that it can provide useful guidance in applications without ground-truth as a validation. This can help assess the fidelity of trained deep neural networks in application scenarios where conditions may be different from the training phase. For example, 3D-SQUIRREL can help assess the 3D-super-resolution fidelity when the raw confocal stack SNR varies (**Fig. 5f, Fig. S11**).

**Fig. 5.**
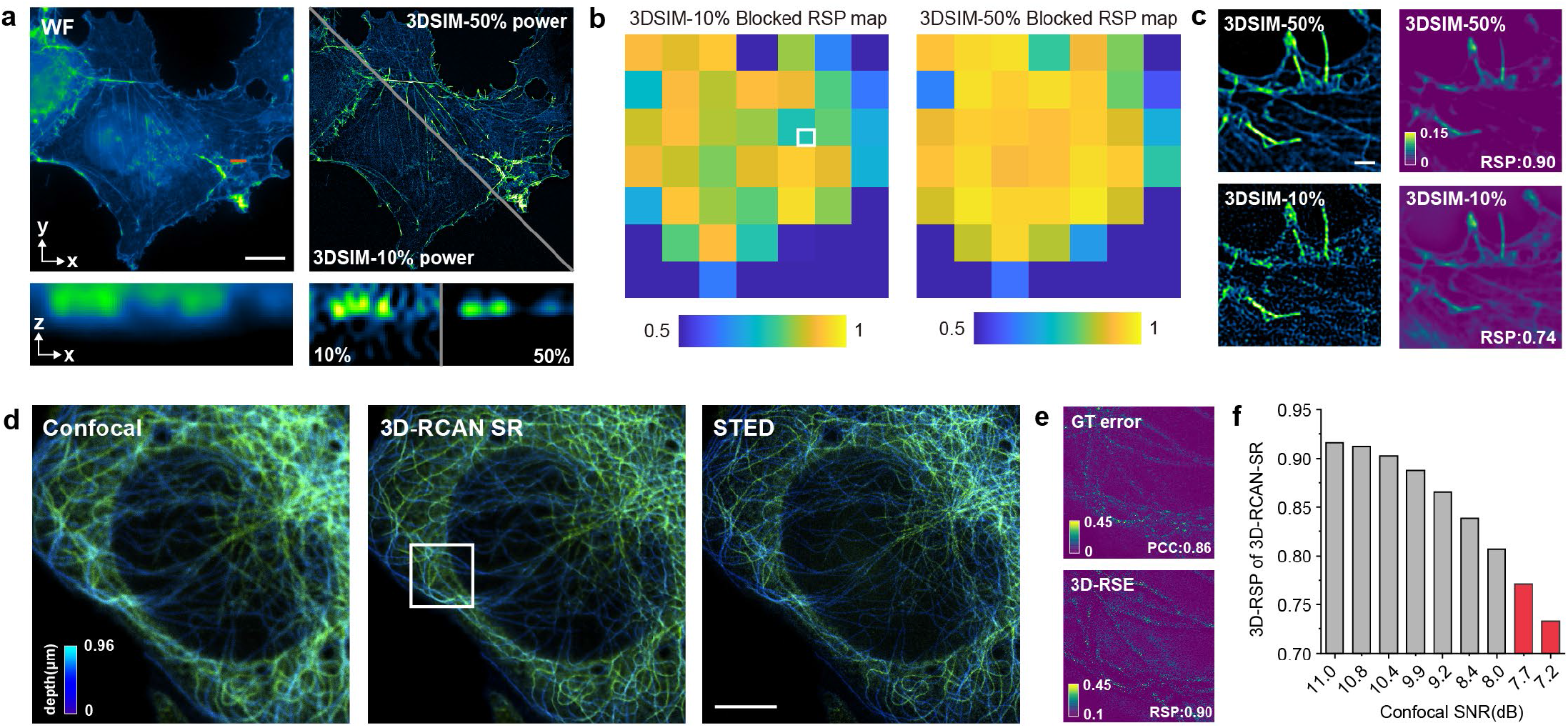
Assessing 3D-super-resolution quality with 3D-SQUIRREL. **a**. Widefield and 3D-SIM images with different illumination power. The xz-slice is made from the orange line. **b**. Blocked 3D-RSP maps of the two 3D-SIM reconstruction results. **c**. Magnified white boxed region in e. **d**. The confocal image stack, and its 3D super-resolution results via 3D-RCAN and STED. **e**. Magnified white boxed region in g. **f**. 3D-RSP of 3D-RCAN super-resolution results with different raw Confocal SNR. Scale bars: 10 *μ*m (a), 1 *μ*m (c), 5 *μ*m (d).

## Discussion

In summary, 3D-ADE is a noise-robust and easy-to-use 3D-deconvolution and evaluation toolkit for volumetric fluorescence imaging (**Fig. S12**), which is applicable to a variety of advanced microscopy techniques. It incorporates robust mathematical models to address noise instability and simplify the cumbersome 3D-PSF calibration process, facilitating broader 3D-deconvolution applications. The proposed 3D-SQUIRREL serves as a criterion to ensure a reliable deconvolution result, and can also be used to improve physical super-resolution microscopy systems and judge the reliability of deep-learning super-resolution results.

Furthermore, for a balanced perspective, the limitations of the tool should be discussed. Although 3D-Ada can automatically estimate the 3D-PSF, interfering factors, such as microscope aberrations, cannot be accounted for in this process. The evaluation results of 3D-SQUIRREL can only serve as a necessary condition for the reliability of super-resolution. As with the limitations of 2D-SQUIRREL, because 3D-SQUIRREL only uses the low-resolution 3D image stack as a reference, it cannot assess small-scale artifacts.

To maximize user accessibility, the code, together with user interactive software, is developed with MATLAB, ImageJ (Fiji) plugin, and an executable file (exe). We also provide a user manual and video guidance (**Video S1**).

## Methods

### Code and software development

The original 3D-Ada deconvolution and 3D-SQUIRREL codes were developed based on MATLAB 2022b version. To facilitate their distribution, we developed their ImageJ plugins based on Java version 1.8. Graphical User Interface (GUI) with an executable file (exe) format is also presented. The code can be further accelerated by a graphics computing unit (GPU). Under the MATLAB testing environment, to perform Ada3D deconvolution on a 1024×1024×40 3D-stack, for 40 iterations, it takes ∼3 minutes for pure CPU computation (Intel i7-12700, 2.10 GHz), and takes only ∼1 minute with GPU acceleration (Nvidia RTX 4080). To evaluate the quality of a 1024×1024×40 3D-stack with a low-resolution reference of the same size, it takes ∼20 seconds for pure CPU computation (Intel i7-12700, 2.10 GHz), and takes only ∼6 seconds with GPU acceleration (Nvidia RTX 4080).

### Self-collected data

This study involves some self-collected data to evaluate the performance of 3D-Ada and 3D-SQUIRREL. We captured the widefield, SD-confocal microscope data using a commercially available NovaSD spinning-disk confocal microscope (Airy Technologies, China). We use a commercially available mouse kidney section (FluoCells Prepared Slide #3, Invitrogen, F24630), a commercial slice of the stem of the *Nerlum* sample (Sagaoptics, China). The image of a Drosophila egg labeled with Tubulin was generously provided by Dr. Guangwei Xin from the Experimental Teaching Center of the School of Life Sciences at Peking University.

### Open-source data

To examine the wide feasibility of the proposed 3D-Ada and 3D-SQUIRREL methods, we collected some open-source data from published papers for validation. The light-field data^7^ of the mouse liver are available at: https://doi.org/10.5281/zenodo.7233421. The two-photon microscope data^5^ of *Pinus radiata* are available at: https://doi.org/10.5281/zenodo.15031504. The EPFL data^22^ used for evaluations are available at: https://bigwww.epfl.ch/deconvolution/deconvolutionlab2. The 4Pi-SIM data^26^ are available at: https://doi.org/10.6084/m9.figshare.25714068. The 4-beam SIM data^25^ are available at: https://zenodo.org/record/6727773. The volumetric confocal-STED paired microtubule datasets^27^ are available at: https://zenodo.org/record/4624364#.YF3jsa9Kibg. The SD-confocal image^15^ of Beta actin is available at: https://downloads.allencell.org/publication-data/label-free-prediction/index.html.The instantSIM data^20^ are available at: https://zenodo.org/record/4624364#.YF3jsa9Kibg.

### Comparisons with Richardson-Lucy-based methods

We compare the performance of 3D-Ada with advanced 3D-Richardson-Lucy-based methods. We tested 3D-RL deconvolution algorithms using both the inner function of MATLAB and that in deconvlab2^22^, which yield similar results. For 3D-EV deconvolution^33^, which is an advanced regularized variant of 3D-RL, we directly run their code at: https://github.com/shengxiao01/Extended-Volume-3D-deconvolution. For blind 3D-RL deconvolution, we initialized the 3D-PSF as a Gaussian PSF with a sigma parameter of 2. The results are generated with 20 iterations.

### Training of 3D-RCAN for super-resolution of confocal stack

To demonstrate the utilization of 3D-SQUIRREL for assessing uncertainties in deep-learning-based super-resolution, we employed an open-source deep-learning dataset of transforming volumetric confocal to STED stacks, using a 3D-RCAN super-resolution neural network. The dataset is publicly available at: https://zenodo.org/record/4624364#.YF3jsa9Kibg. We employed a 3D-RCAN model with 3 residual blocks and 5 residual groups, as those were used for training on this dataset^27^. Other hyperparameters, including learning rate, patch size, and epochs, were chosen as they were used on this dataset. The training was performed on a local workstation with an NVIDIA RTX 4080 graphics computing unit (GPU).

### Image quality metrics

For cases with known ground-truth, we employ PSNR and SSIM to evaluate the image quality metrics. Given a ground-truth image stack *g* and a recovered image stack *f*, the PSNR value is calculated as follows:

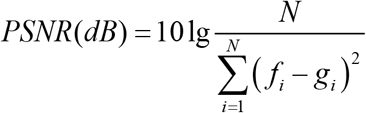

The SSIM value is calculated as:

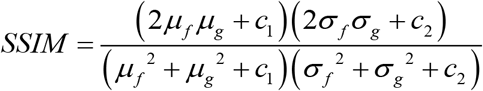

where *μ*_*f*_ and *μ*_*g*_ are their average value, *σ*_*f*,_ and *σ*_*g*_ are their standard deviation value, *σ*_*f*_ *σ*_*g*_ is the covariance of *f* and *g, c*_1_ and *c*_2_ are two constants to stabilize the result (*c*_1_=(*k*_1_*L*)^2^, *c*_2_=(*k*_2_*L*)^2^, *L* is the dynamic range of pixel value, *k*_1_=0.01, *k*_2_=0.03).

### 3D-data visualization

To visualize the 3D data in a finer way, we adopted pseudo-color projection and 3D rendering with Imaris Viewer software.

## Author contribution

Y.H. and P.X. conceived the idea of 3D-Ada deconvolution and 3D-SQUIRREL assessment of 3D super-resolution errors. P.X. and M.L. supervised this project. Y. H. developed the mathematical framework, wrote all the code, and composed all the figures. W.W. provided the SD-confocal and 3D-SIM data. R.C. provided the 2D-SIM data. Y.F., X.S., and D.K. provided valuable suggestions in arranging the paper. Y. H., P. X., and M. L. wrote the manuscript with inputs from all the authors. All authors discussed the results presented in the paper.

## Acknowledgements

This work was supported by the National Natural Science Foundation of China (62335008, 62405010 to M. L.; 62025501 to P. X.), National Key R&D Program of China (2022YFC3401100 to P. X.), and Major Basic Research Project of the Natural Science Foundation of Shandong Province (ZR2024ZD27 to P. X.). We thank National Center for Protein Sciences at Peking University in Beijing, China, for assistance with super-resolution imaging.

