## Supplementary Information for "Data-adaptive three-dimensional deconvolution and evaluation for volumetric fluorescence microscopy"

### Note S1. 3D-Ada noise-suppression model

Here, we develop the adaptive noise control model for volumetric fluorescence imaging on the basis of 3D-framelet. The 3D framelet transform is a redundant transform that can better capture key edge information over Fourier or spatial derivative-based methods. Denote the 1D framelet bases as $\phi$, $\psi_1$, and $\psi_2$. The 3D framelet base can be generated by the tensor product of the 1D framelet base:

$$\Phi(x,y,z) = \phi(x) \otimes \phi(y) \otimes \phi(z) \tag{1-1}$$

$$\begin{aligned}\{\Psi(x,y,z)\} = \{&\phi(x)\otimes\psi_{l_1}(y)\otimes\psi_{l_2}(z), \phi(x)\otimes\phi(y)\otimes\psi_{l_2}(z),\\ &\phi(x)\otimes\psi_{l_1}(x)\otimes\phi(z), \psi_{l_1}(x)\otimes\phi(y)\otimes\psi_{l_2}(z),\\ &\psi_{l_1}(x)\otimes\phi(y)\otimes\phi(z), \psi_{l_1}(x)\otimes\psi_{l_2}(y)\otimes\phi(z), 1 \le l_1, l_2 \le 2\}\end{aligned} \tag{1-2}$$

where $\Phi$ $(x, y, z)$ is the scaling function, $\Psi$ $(x, y, z)$ is the wavelet function. The framelet coefficient can be obtained by the inner product of the 3D data with the 3D framelet bases.

The denoising of 3D framelet can be achieved by soft-thresholding its decomposition coefficients:

$$f = F_{3D}^{-1}\mathcal{T}_\lambda(F_{3D}g) \tag{1-3}$$

where $F_{3D}$ denotes the 3D framelet transform, $F_{3D}^{-1}$ denotes the inverse framelet transform.

However, a key issue is that this thresholding is uniform. For instance, we inspect the average energy of the 2D-framelet and 3D-framelet spectra of a typical volumetric image stack (**Fig. S1a-c**). It can be found that the 3D-framelet spectra are much more uneven, with more energy concentrated in the low-frequency parts compared with the 2D-framelet spectra. This indicates that some 3D structures, despite being our target signals, may not be sparsely represented by 3D-framelet, due to the anisotropic 3D fluorescence microscopic imaging process. As a result, when uniformly thresholding the 3D-framelet spectra, there is a great hazard that some desired high-frequency information would be filtered.

To address this issue and preserve the high-frequency details when thresholding the 3D-framelet spectra, we employ Stein's unbiased estimate of risk (SURE) to analyze each subband of the 3D-framelet spectra. Suppose $c$ is the collection of the coefficients in a certain subband with $d$ elements, and $t$ is the thresholding value chosen for the subband. The SURE method assumes an estimated error with a normal distribution. The resulting optimization problem to minimize the risk is as follows:

$$t_{SURE} = \arg\min_{t} d - 2 \cdot \#\{i : |c_i| \le t\} + \sum_{i=1}^{d} \left[\min(c_i, t)\right]^2 \tag{1-5}$$

where #{} denotes the number of elements that satisfy the condition within the braces, $t_{SURE}$ is the thresholding value decided by the SURE criterion.

Despite SURE shrinkage decision could bring near optimal value in respect of the mean-square-error (MSE) measurement, the effect of the forward blurring operator in the inverse problem should also be considered. More frequently, we found that SURE shrinkage would give an overconfident estimation of the thresholding value and smear the desired high-frequency component. Therefore, we reformulate a subband-weighted thresholding denoising method, using the relative ratio of the SURE-decided thresholding value as the weights:

$$w^{l,i,j,k} = t_{SURE}{}^{l,i,j,k} / t_{SURE}{}^{1,1,1,1} \quad i, j, k \in \{1, 2, 3\} \tag{1-6}$$

$$\mathcal{T}_{w^{l,i,j,k},\lambda}\left(c^{l,i,j,k}\right) = \operatorname{sgn}\left(c^{l,i,j,k}\right) \cdot \max\left(0, c^{l,i,j,k} - w^{l,i,j,k} \cdot \lambda\right) \tag{1-7}$$

where the indices *i,j,k* denote the framelet filter that is applied for the *x,y,z* dimensions, $l$ denotes the decomposition level, $\lambda$ denotes the applied soft-thresholding value, $c$ denotes the framelet coefficients.

To demonstrate the effectiveness of this subband-weighted 3D-framelet thresholding method, we have conducted a simulation denoising experiment with known ground-truth (**Fig. S1d**). While applying the same thresholding to the subband representing the lowest frequency features, the subband-weighted 3D-framelet thresholding shows evidently finer preservation of high-frequency details. The quantitative analysis results indicate that the subband-weighted thresholding method could bring a higher PSNR value (**Fig. S1e**). When achieving the respective optimal PSNR value, the subband-weighted thresholding method preserves the details better as reflected in FWHM analysis and the Fourier spectrum (**Fig. S1f, g**). We hereafter examine the performance of this data-adaptive noise reduction method in real biological samples. The results show that the proposed method demonstrates advantages over spatial derivative-based methods in both visual perception and quality metrics (**Figs. S2, 3**).

### Note S2. 3D-Ada automatic 3D-PSF estimation

- **Main concept**

Calibration of 3D-PSF is always a cumbersome issue for 3D-deconvolution. Here, we develop an automatic 3D-PSF estimation method. The rationale is that essentially any PSF has a key parameter that controls its FWHM, which decides the blurry condition (i.e., resolution). By leveraging some prior knowledge of the microscopic imaging process (mainly the Fourier spectra are compactly supported with a circular shape, and signals show correlations), some advanced Fourier ring correlation (FRC) methods, such as the decorrelation analysis, can estimate the imaging resolution with acquired images. Then, once the resolution estimation is accurate, we can reliably infer the 3D-PSF. The 3D-Ada noise model also aids the feasibility of this approach in practical imaging, where complex noise is present.

- **Robust resolution analysis with a multi-filtering strategy**

Here, we utilize the decorrelation analysis to estimate the resolution, which is a widely adopted, advanced single-image FRC-derived method.

We first calculate the resolution of the lateral image through decorrelation analysis. Given a 3D stack $f_{stack}$, we first obtain its average intensity projection (AIP) $f_{AIP}$, which can mitigate the noise, then its decorrelation curve is calculated as:

$$d(r)=\frac{\int \mathrm{Re}\left\{ f_{AIP}\left(\vec{k}\right) f_{AIPn}^{*}\left(\vec{k}\right) M\left(\vec{k};r\right)\right\} dk_x dk_y}{\sqrt{\int \left| f_{AIP}\left(\vec{k}\right)\right|^2 dk_x dk_y \int \left| f_{AIPn}\left(\vec{k}\right) M\left(\vec{k};r\right)\right|^2 dk_x dk_y}} \tag{2-1}$$

where $\vec{k}=[k_x,k_y]$ is the frequency-domain coordinate, the subscript n denotes a normalization operation, $M(\vec{k};r)$ is the binary mask with a radius of $r$. The resolution then can be determined by filtering the original image with multiple Gaussian low-pass filters until there is no critical point in the curve.

Since the sampling in the axial dimension is usually much lower than that in the lateral dimension, it is usually hard to precisely determine the axial resolution through the decorrelation analysis. Therefore, we adopt a detour strategy that inspects the Fourier spectrum of the x-z slice and infers the axial resolution through the x-z feature obtained in the Fourier spectrum.

To achieve so, we first obtain an average intensity projection (AIP) x-z slice of the 3D-stack, denoted by fxz, which is up-sampled using the bilinear method to ensure that the pixels in x and z are the same. Then we perform a removal of the DC component:

$$f_{xz} = f_{xz} - Avg\left(f_{xz}\right) \tag{2-2}$$

Then 2D Hanning window is applied to suppresses spectral leakage of Fourier transform:

$$\begin{cases} f_{xz} = f_{xz} \cdot W(x,z) \\ W(x,z) = W_x(x) \cdot W_z(z) \\ W_x(x) = 0.5\left[1 - \cos\left(\dfrac{2\pi x}{d}\right)\right] \\ W_z(z) = 0.5\left[1 - \cos\left(\dfrac{2\pi z}{d}\right)\right] \end{cases} \tag{2-3}$$

Then perform the Fourier transform of fxz, and obtain its normalized power spectrum density, denoted as $PSD_{2D}(u,v)$. Then subtract the lateral and axial 1D intensity as follows:

$$\begin{cases} PSD_x(u) = \dfrac{1}{2 \cdot border + 1} \sum_{v=-border}^{border} PSD_{2D}(u,v) \\ PSD_z(v) = \dfrac{1}{2 \cdot border + 1} \sum_{u=-border}^{border} PSD_{2D}(u,v) \end{cases} \tag{2-4}$$

where we select *border* as 11.

Then each 1d PSF can be fitted by a power function as follow:

$$PSD(f) = A \cdot f^{-\alpha} + B \tag{2-5}$$

where $A$ is the amplitude coefficient, α is the attenuation coefficient, and $B$ is the noise base value.

The noise base value can be estimated via the high-frequency component average value:

$$\begin{cases} B_x = Avg\left(PSD_x(u) \middle| u > 0.8 * \max(u)\right) \\ B_z = Avg\left(PSD_z(v) \middle| v > 0.8 * \max(v)\right) \end{cases} \tag{2-6}$$

Then transform the power function to a linear format as:

$$\ln\left(PSF(f) - B\right) = \ln(A) - \alpha \cdot \ln(f) \tag{2-7}$$

Then the value of A and α can be obtained by simple least-square-fitting.

Then we find the cut-off frequency of the lateral and axial direction respectively

$$\begin{cases} u_{cx} = \min\left\{ u \middle| A_x \cdot u^{-\alpha_x} + B_x < \dfrac{\max(PSD_x)}{e} \right\} \\ v_{cz} = \min\left\{ v \middle| A_z \cdot v^{-\alpha_z} + B_z < \dfrac{\max(PSD_z)}{e} \right\} \end{cases} \tag{2-8}$$

Then the axial resolution is estimated using the lateral decorrelation result:

$$d_{res\_z} = d_{res\_xy} \cdot \frac{v_{cz}}{u_{cx}} \tag{2-9}$$

However, noise in practical fluorescence imaging conditions would mislead the decorrelation analysis, causing inaccurate resolution estimation (**Fig. S4**). We solve this issue by applying 3D-Ada-based denoising before performing the decorrelation analysis.

Selection of the thresholding value is a tricky issue for different samples and imaging conditions. A large thresholding weight would smear some high-frequency details, causing a lower resolution estimated value. While a small thresholding weight would leave residual noise, it would also influence the decorrelation analysis result. To solve this issue, we develop a multi-filtering strategy, which is pre-filtering the image stack with multiple thresholding values:

$$\begin{cases} f^i = F_{3D}^{-1} \mathcal{T}_{\lambda^i}\left(F_{3D} f_{stack}\right) \\ \lambda = \{0.01, 0.005, 0.001, 0\} \end{cases} \tag{2-10}$$

The definitions of the symbols in (2-4) are detailed in Supplementary Note 1. Then the resolution is designated with $f_{MIP}^i$ with the highest decorrelation cut-off frequency:

$$k_c = \max\left\{decorr\left(f_{AIP}^i\right)\right\} \tag{2-11}$$

$$d_{res\_xy} = \frac{2}{k_c} \tag{2-12}$$

Significance of this multi-filtering strategy in assessing resolution in practical noisy conditions is shown in **Fig. S4**.

- **Translational function**

Another key part of the conceived automatic 3D-PSF estimation pipeline is deducing key parameters of 3D-PSF using the estimated resolution value. Below, we first deduce this process using a Gaussian PSF as an example.

Suppose two adjacent points in an image, after a Gaussian-shaped PSF blurring, exhibit the following intensity profiles along one dimension:

$$I(x) = e^{-\frac{x^2}{2\sigma^2}} + e^{-\frac{(x-d)^2}{2\sigma^2}} \tag{2-13}$$

where $x$ represents the spatial coordinate, and $\sigma$ represents the standard deviation of the Gaussian kernel.

At the midpoint, we have the following intensity:

$$I(\frac{d}{2}) = 2e^{-\frac{d^2}{8\sigma^2}} \quad (2\text{-}14)$$

The resolution criterion can be interpreted as when the middle point of the intensity profile reaches some critical value:

$$I(\frac{d}{2}) = 2e^{-\frac{d^2}{8\sigma^2}} = I_{critical} \quad (2\text{-}15)$$

where $I_{critical}$ is the criterion whether the two points can be separated (i.e., resolution).

Then it can be determined that the $\sigma$ parameter of the Gaussian PSF has the following relationship with the resolution value, which we termed a translational function:

$$d_{res} = \sigma\sqrt{-8ln(\frac{I_{critical}}{2})} \quad (2\text{-}16)$$

Gathering the above discussions, taking a Gaussian 3D-PSF as an example, the image-drive automatic 3D-PSF calibration can be achieved as follows:

$$\begin{cases} \sigma_{xy} = \dfrac{d_{res\_xy}}{\sqrt{-8ln(\frac{I_{critical}}{2})}} \\ \sigma_z = \dfrac{d_{res\_z}}{\sqrt{-8ln(\frac{I_{critical}}{2})}} \\ PSF_{3D} = \exp\left[-\left(\dfrac{x^2+y^2}{2\cdot\sigma_{xy}^2} + \dfrac{z^2}{2\cdot\sigma_z^2}\right)\right] \end{cases} \quad (2\text{-}17)$$

Some empirical criteria were used to decide the value of $I_{critical}$ as the resolution criterion, such as the Rayleigh, Abbe, and Sparrow criteria. However, the practical FRC method to estimate the resolution value is essentially a measurement of signal-to-noise ratio, and is not directly related to the above-mentioned empirical criteria. For an empirical usage, we set $I_{critical} = 0.5$ and find that in most cases it corresponds well with the resolution value obtained by decorrelation analysis. It should be mentioned that the choice of $I_{critical}$ does not influence the physical proportional relationship of $d_{res}$ and $\sigma$.

- **Generalization to any 3D-PSF shapes**

Above, we present our automatic 3D-PSF estimation pipeline using a Gaussian shape as an example. This method can essentially be generalized to any 3D-PSF shapes, which may involve some sidelobe calculations to better approximate the imaging process. This process can be easily achieved by matching the lateral and axial full-width-at-half-maxima (FWHM) of the 3D-PSF with a calculated Gaussian PSF. It can be easily deduced that the relationship of the FWHM and σ parameters in a Gaussian function is as follows:

$$FWHM = 2\sqrt{2\ln 2}\cdot\sigma \tag{2-18}$$

For a 3D-PSF $PSF^{\gamma_{xy},\gamma_z}(x,y,z)$ with lateral feature parameterized by $\gamma_{xy}$ and axial feature parameterized by $\gamma_z$, it can be reconstructed by the following formula:

$$\underset{\gamma_{xy}}{\arg\min}\left\| FWHM_{xy}\left\{PSF^{\gamma_{xy},\gamma_z}(x,y,z_{center})\right\} - 2\sqrt{2\ln 2}\cdot\sigma_{xy}\right\|_2 \tag{2-19}$$

$$\underset{\gamma_z}{\arg\min}\left\| FWHM_z\left\{PSF^{\gamma_{xy},\gamma_z}(x_{center},y_{center},z)\right\} - 2\sqrt{2\ln 2}\cdot\sigma_z\right\|_2 \tag{2-20}$$

In cases $PSF^{\gamma_{xy},\gamma_z}(x,y,z_{center})$ has an explicit expression, its parameters $\gamma_{xy}$ and $\gamma_z$ would have analytical solutions. Otherwise optimization method can be adopted to search $\gamma_{xy}$ and $\gamma_z$ with some appropriately defined boundary conditions.

For example, consider a Gaussian-Lorentzian 3D-PSF that can approximate the widefield volumetric imaging conditions:

$$PSF(x,y,z) = \frac{2\pi\cdot NA^2\cdot(n/\lambda)^2}{1+\left(\pi\cdot NA^2\cdot n/\lambda\cdot z\right)^2}\cdot\exp\left[-\frac{2\pi^2\cdot NA^2\cdot(n/\lambda)^2\cdot\left(x^2+y^2\right)}{1+\left(\pi\cdot NA^2\cdot n/\lambda\cdot z\right)^2}\right] \tag{2-21}$$

where $NA$ is the numerical aperture, $n$ is the refractive index, and $\lambda$ is the imaging wavelength. If using imaging parameters to reconstruct this parameter, the lateral and axial pixel size should also be provided to change the real distance to pixel count. Despite a more complex model using more imaging parameters to reconstruct this PSF, essentially, these parameters decide scaling factors $\gamma_{xy}$ and $\gamma_z$ that control the lateral and axial FWHM of this PSF.

$$PSF^{\gamma_{xy},\gamma_z}(x,y,z) = \frac{1}{1+\gamma_z\cdot z^2}\cdot\exp\left[-\frac{\gamma_{xy}\cdot\left(x^2+y^2\right)}{1+\gamma_z\cdot z^2}\right] \tag{2-22}$$

Using the formulas (2-19) and (2-20), it can be deduced that once we obtain the Gaussian PSF parameters, the PSF described in (2-22) can be calculated as:

$$\begin{cases}\gamma_{xy} = \dfrac{1}{\sigma_{xy}}\\[2ex] \gamma_z = \dfrac{1}{\ln 2\cdot\sigma_z}\end{cases} \tag{2-23}$$

3D-PSF with other shapes can be reconstructed similarly. For ease of usage in biologists and microscopists, here in this study, we use a Gaussian-Lorentzian 3D-PSF for a widefield detection system, and a Gaussian 3D-PSF for a point-detected system, since previous literature points out that they have a well Gaussian approximation.

#### Table S1. Process of automatic 3D-PSF estimation

**Execution of 3D-Ada automatic 3D-PSF estimation**

**Step0. Input:** Image stack $f_{stack}$, detection type (widefield or point-detected)

**Step1. Multi-filtering of the image stack**

$$f^{i}=F_{3D}^{-1}\mathcal{T}_{\lambda^{i}}\left(F_{3D}f_{stack}\right);\ \ \lambda=\left\{0.01,0.005,0.001,0\right\}$$

**Step2. Decorrelation analysis**

$$d\left(r\right)=\frac{\int\mathrm{Re}\left\{f_{AIP}\left(\vec{k}\right)f_{AIPn}^{*}\left(\vec{k}\right)M\left(\vec{k};r\right)\right\}dk_{x}dk_{y}}{\sqrt{\int\left|f_{AIP}\left(\vec{k}\right)\right|^{2}dk_{x}dk_{y}\int\left|f_{AIPn}\left(\vec{k}\right)M\left(\vec{k};r\right)\right|^{2}dk_{x}dk_{y}}}$$

$$k_{c}=\max\left\{decorr\left(f_{AIP}^{i}\right)\right\}\qquad d_{res\_xy}=\frac{2}{k_{c}}$$

**Step3. Anisotropy analysis**

$$d_{res\_z}=d_{res\_xy}\cdot\frac{v_{cz}}{u_{cx}}$$

**Step4. Applying translational function**

$$d_{res\_xy}=\sigma_{xy}\sqrt{-8ln(\frac{I_{critical}}{2})}\qquad d_{res\_z}=\sigma_{z}\sqrt{-8ln(\frac{I_{critical}}{2})}$$

**Step5. Parameter matching and constructing 3D-PSF**

$$\underset{\gamma_{xy}}{\arg\min}\left\|FWHM_{xy}\left\{PSF^{\gamma_{xy},\gamma_{z}}(x,y,z_{center})\right\}-2\sqrt{2\ln 2}\cdot\sigma_{xy}\right\|_{2}$$

$$\underset{\gamma_{z}}{\arg\min}\left\|FWHM_{z}\left\{PSF^{\gamma_{xy},\gamma_{z}}(x_{center},y_{center},z)\right\}-2\sqrt{2\ln 2}\cdot\sigma_{z}\right\|_{2}$$

$$3D-PSF=PSF^{\gamma_{xy},\gamma_{z}}\left(x,y,z\right)$$

### Note S3. 3D-FISTA inverse problem solver

FISTA iteration has shown great efficacy in solving the 2D convex non-smooth inverse problem and is widely adopted. It mainly solves the optimization problem with the following formulation:

$$\min_{f} \; F(f) \equiv F_1(f) + F_2(f) \tag{3-1}$$

where $F(f)$ is the total cost function, $F_1(f)$ is a convex and smooth function, and $F_2(f)$ is a convex but possibly unsmooth function.

In the proposed 3D-Ada deconvolution framework, we have:

$$F_1(f) = \left\| g_{3D} - A_{3D} \cdot f_{3D} \right\|_2^2 \tag{3-2}$$

$$F_2(f) = \lambda \sum_{l,i,j,k} w_{l,i,j,k} \left\| W_{l,i,j,k} f_{3D} \right\|_1 \tag{3-3}$$

where $g_{3D}$ denotes the recorded volumetric images by a microscope, $g_{3D}$ is the image to be recovered, $A_{3D}$ is the matrix form of the 3D-PSF, $\lambda$ is the regularization parameter, w is the weight decided by the SURE criterion, *i,j,k* denote the framelet filter that is applied for the *x,y,z* dimensions, *l* denotes the decomposition level.

The gradient method that can minimize $F_1(f)$ is:

$$f_{3D}{}^{k} = f_{3D}{}^{k-1} - t_k \nabla F_1\left( f_{3D}{}^{k-1} \right) \tag{3-4}$$

Soft-thresholding can be utilized to minimize $F_2(f)$:

$$\mathcal{T}_{w,\lambda}(x) = \left( |x| - w \cdot \lambda \right) \operatorname{sgn}(x) \tag{3-5}$$

where $x$ is the coefficient to be soft-thresholded.

We derive the 3D version of FISTA iteration to solve the total minimization problem, as listed in **Table S3**.

**Table S2. Execution process of 3D-Ada deconvolution**

**Execution of 3D-Ada deconvolution**

**Step0. Input:** Degenerated image stack $g_{3D}$, system PSF $PSF_{3D}$, regularization parameter $\lambda$

**Step1. Calculation of SURE-based thresholding weight:**

$$t_{SURE} = \arg\min_{t} d - 2 \cdot \#\left\{ i : \left| c_i \right| \le t \right\} + \sum_{i=1}^{d} \left[ \min\left( c_i, t \right) \right]^2$$

$$w^{l,i,j,k} = t_{SURE}{}^{l,i,j,k} / t_{SURE}{}^{1,1,1,1} \quad i, j, k \in \{1, 2, 3\}$$

**Step2. 3D-Ada deconvolution iteration**

Initialization: Let $f_{3D}{}^{0} = g_{3D}$, $t_0 = 1$, L is the Lipschitz constant of the system PSF.

Iteration:

The $k$th iteration ($k \ge 1$):

$$f_{3D}{}^{k} = f_{3D}{}^{k-1} - t_k \nabla F_1\left( f_{3D}{}^{k-1} \right)$$

$$f_{3D}{}^{k}{}' = W^T \mathcal{T}_{w \cdot \lambda / L}\left( W\, f_{3D}{}^{k} \right)$$

$$f_{3D}{}^{k} = f_{3D}{}^{k}{}'$$

$$t_{k+1} = \frac{1 + \sqrt{1 + 4t_k^2}}{2}$$

$$f_{3D}{}^{k+1} = f_{3D}{}^{k} + \left( \frac{t_k - 1}{t_{k+1}} \right)\left( f_{3D}{}^{k} - f_{3D}{}^{k-1} \right)$$

Until maximum iteration time reaches

**Table S3.** Hyperparameters of the deconvolution applied in the study

| Data number | Detection type | Regularization | Iteration |
|---|---|---|---|
| Fig. 2b | Widefield | 0.0001 | 100 |
| Fig. 2d | Widefield | 0.0001 | 400 |
| Fig. 2e | Widefield | 0.0005 | 100 |
| Fig. 3a | Widefield | 0.0001 | 60 |
| Fig. 3b | Widefield | 0.0001 | 40 |
| Fig. 3c | Point-detection | 0.0005 | 100 |
| Fig. 3d | Point-detection | 0.0005 | 100 |
| Fig. 3e | Point-detection | 0.001 | 60 |
| Fig. 3f | Widefield | 0.0001 | 60 |

### Note S4. Development of 3D-SQUIRREL and its interpretation in the 3D-deconvolution process

- **3D-SQUIRREL optimization problem statement**

The SQUIRREL (super-resolution quantitative image rating and reporting of error locations) algorithm was first proposed by Culley et al. to quantify the possible artifacts in the super-resolution process. The 2D-SQUIRREL tool has been widely applied to a variety of scenarios as an important standard to quantify super-resolution quality. However, to assess the volumetric imaging case, it ought to be modified since the 2D calculation discards the 3D structural causality. Here, we proposed its 3D version, termed 3D-SQUIRREL.

We develop the calculation procedure of 3D-SQUIRREL following a similar step to that which was done for 2D-SQUIRREL calculation. Given a resolution-limited volumetric stack $I_D$ and a resolution-improved stack $I_S$, we first correct the shift of $I_S$ with respect to $I_D$.

$$I_{ST}(x,y,z) = I_S(x-\Delta x, y-\Delta y, z-\Delta z) \tag{4-1}$$

It is allowed to apply a linear intensity transformation of $I_{ST}$ to allow it to match $I_D$ finer:

$$I_{ST\gamma}(\alpha,\beta) = \alpha \cdot I_{ST} + \beta \tag{4-2}$$

We define the 3D resolution scaling function (RSF) shall have a shape that is similar to the PSF. In the 2D-RSF implementation, a Gaussian function is applied. While for our 3D imaging cases, we employ a Gaussian-Lorentz shape for the widefield system, and a 3D-Gaussian shape for the point-detected system. The resulting joint optimization problem is as follows:

$$\underset{\alpha,\beta,\sigma_{xy},\sigma_z}{\arg\min} \left\| I_D - I_{ST\gamma}(\alpha,\beta) \otimes RSF(\sigma_{xy},\sigma_z) \right\|_2 \tag{4-3}$$

- **Two-stage genetic algorithm for solving the optimization problem**

The optimization problem can in principle be solved by a heuristic algorithm to solve this optimization problem, which was also adopted in the 2D-SQUIRREL algorithm. However, as the problem progresses into 3D, more optimization parameters and voxels would pose difficulty in solving this problem. Heuristic algorithms are random and have the hazard of getting trapped in a local optimum that may be far from the global optimal value. We adopted a genetic algorithm (GA) to solve (4-3), but found that the solution turns out to be rather unstable with different tests, and in most cases, they fail to find a near-optimal solution.

To solve this issue, we developed a novel optimization strategy to stabilize this process. Considering that the Pearson correlation coefficient (PCC) is an indicator irrelevant of the absolute intensity difference, we first search for the RSF parameter and ignore $\alpha$ and $\beta$. For simplicity of discussion, denote:

$$I_{ST\gamma}{}^{\sigma_{xy},\sigma_z} = I_{ST\gamma}\left(\alpha,\beta\right)\otimes RSF\left(\sigma_{xy},\sigma_z\right) \tag{4-4}$$

Then the first-stage GA searching problem becomes:

$$\underset{\sigma_{xy},\sigma_z}{\arg\min}\ \text{1-PCC}\left\{I_D, I_{ST\gamma}{}^{\sigma_{xy},\sigma_z}\right\} \tag{4-5}$$

The second-stage optimization problem becomes:

$$\underset{\alpha,\beta}{\arg\min}\left\|I_D - I_{ST\gamma}{}^{\sigma_{xy},\sigma_z}\right\|_2 \tag{4-6}$$

which can be directly solved by applying an affine normalization.

**Table S4.** Evaluation of the deconvolution results of real samples shown in the study

| Data number | 3D-RSP |
|---|---|
| Fig. 3a | 0.85 |
| Fig. 3b | 0.80 |
| Fig. 3c | 0.91 |
| Fig. 3d | 0.99 |
| Fig. 3e | 0.93 |
| Fig. 3f | 0.81 |

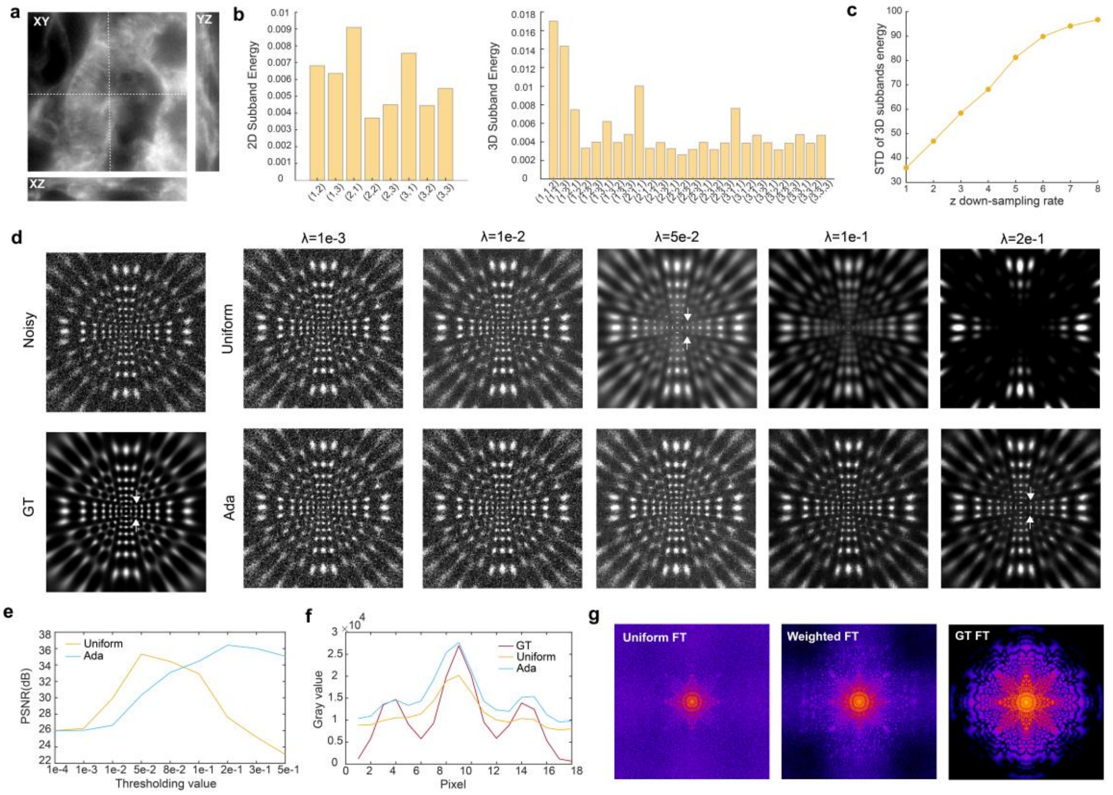


**Fig. S1. Motivation and effectiveness of the proposed adaptive noise model for 3D data. a.** A 3D volumetric microscopic image of a mouse kidney section cell. **b.** Inspecting the features of the 3D-data's 2D-Framelet coefficients (performed on its central plane) and 3D-Framelet coefficients. The ($i$, $j$, $k$) denotes the applied filter number (1: low-pass filter; 2 and 3: high-pass filters) for $x$, $y$, and $z$ dimensions. The 3D coefficients in different sub-bands show larger variance. **c.** The variance in each 3D-Framelet sub-bands shows larger variance with different axial sampling rates. These observations suggest the necessity of an adaptive thresholding model for noise reduction in 3D. **d.** Denoising performance of uniform thresholding and data-adaptive thresholding of the 3D-Framelet spectra. **e.** PSNR metrics of the denoised images via uniform thresholding and data-adaptive thresholding of the 3D-Framelet spectra. **f.** The intensity profiles along the two arrowheads shown in f. **g.** Fourier spectra of the optimal denoised images through uniform thresholding and sub-band-weighted thresholding of the 3D-Framelet spectrum.

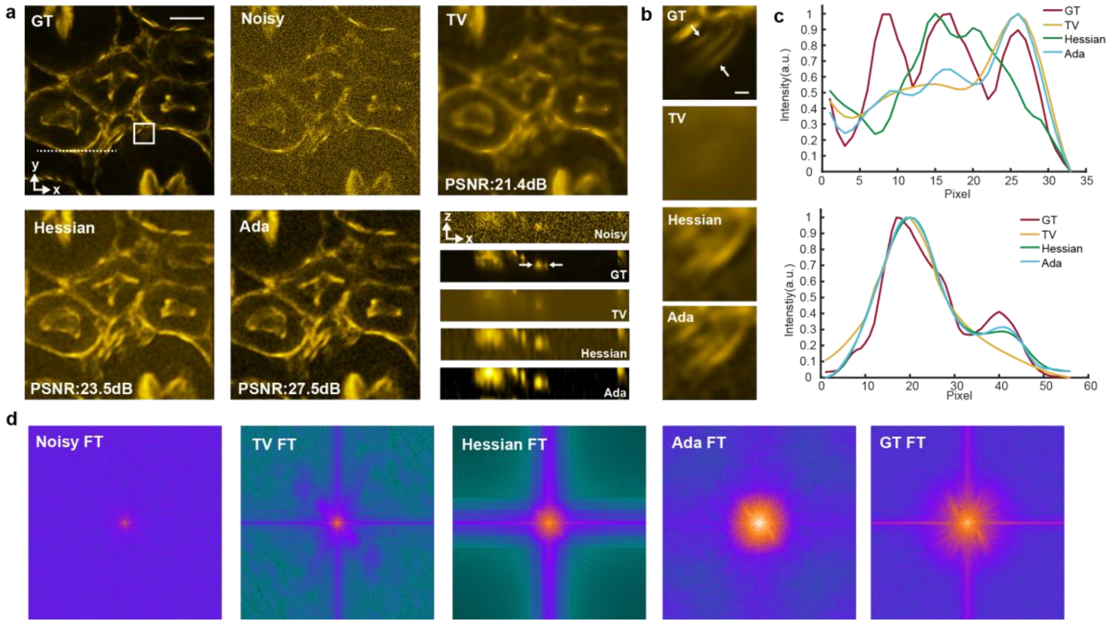


**Fig. S2. Comparison of the noise suppression performance of different noise models for volumetric imaging. a.** Denoising performance of TV, Hessian, and the proposed Ada method in a mouse kidney cell sample labeled by phalloidin. The ground-truth image is captured using a spinning-disk confocal microscope. Optimal parameters of TV, Hessian, and Ada are searched in Fig. S3. The results here are obtained with parameters that produced the highest fidelity metrics. **b.** Magnified white boxed region in a. **c.** Intensity profiles along the two arrowheads in a and b, respectively. d. The Fourier spectra of different results show that Ada finely preserves the high-frequency information, while the gradient-based methods lose these critical components. Scale bars: 10 $\mu$m (a), 1 $\mu$m (b).

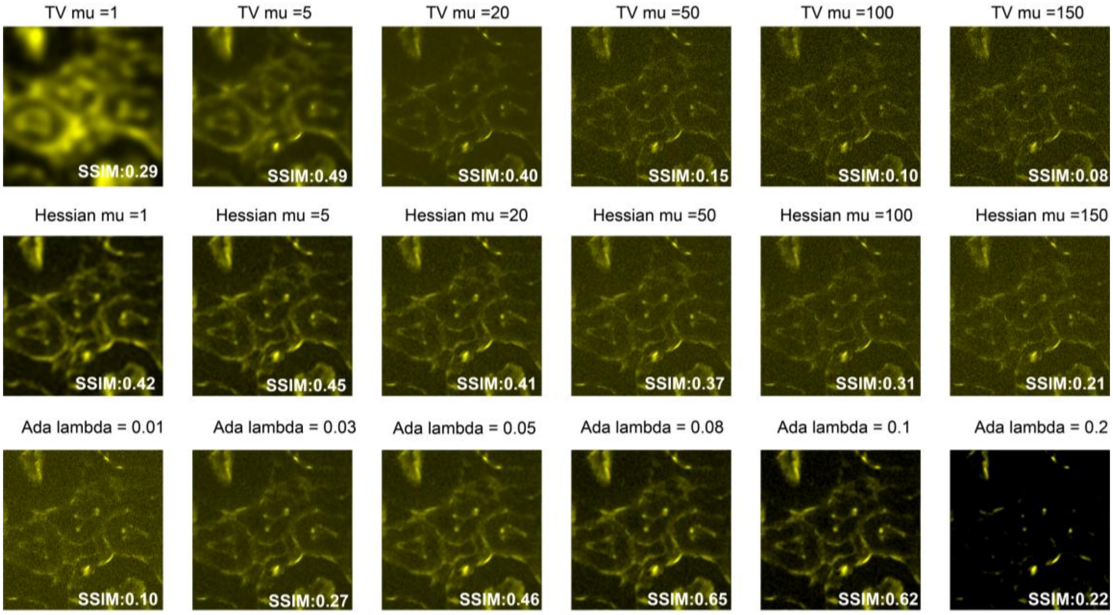


**Fig. S3. Comparison of the noise reduction performance of different noise models under different parameters.**

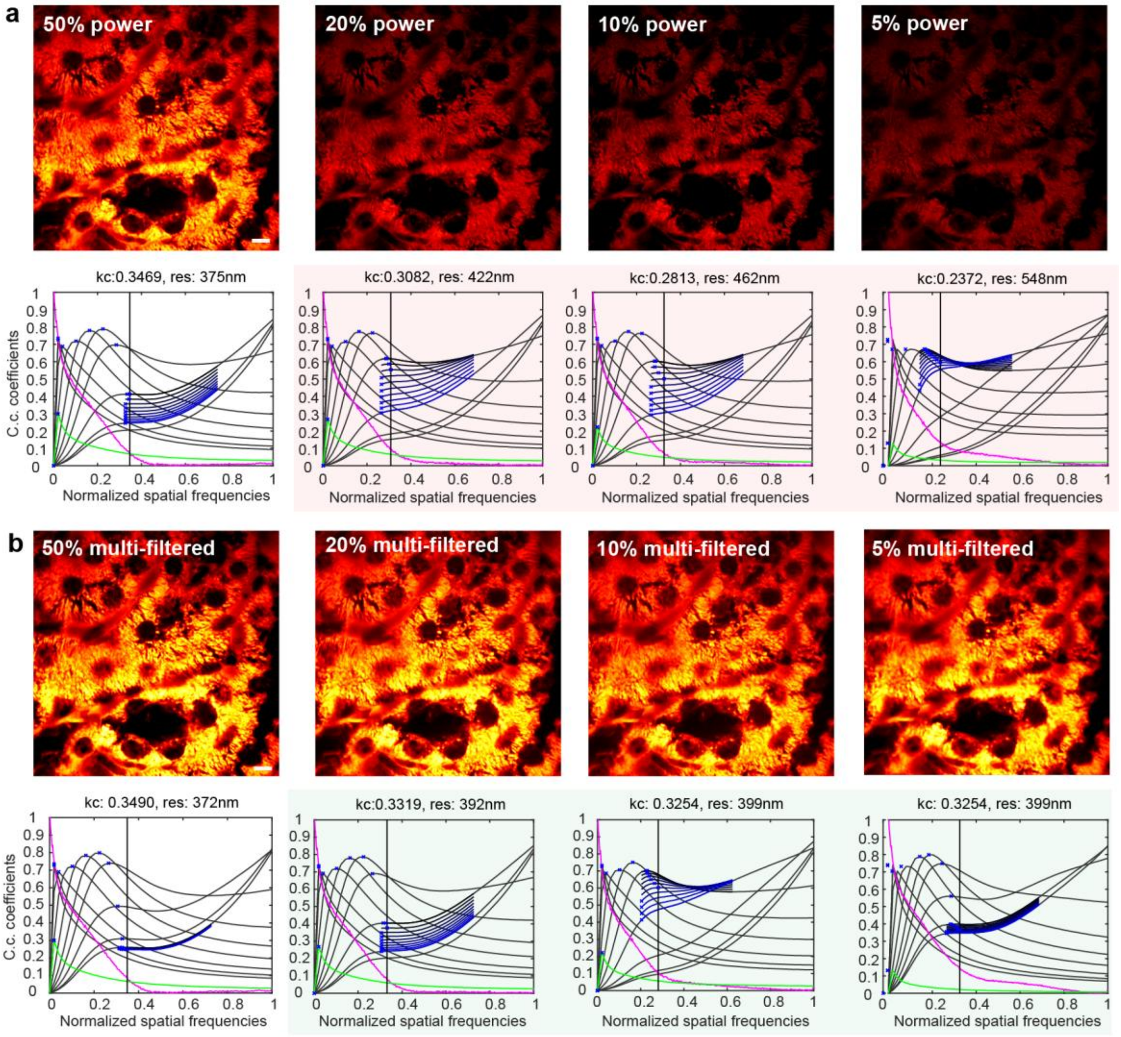


**Fig. S4. Demonstrations of the multi-filtering strategies to improve the noise robustness of automatic 3D-PSF estimation. a.** Raw captured SD-confocal images with gradient illumination power, and the decorrelation analysis results. **b.** Multi-filtered SD-confocal images and the decorrelation analysis results. Scale bars: 10 $\mu$m.

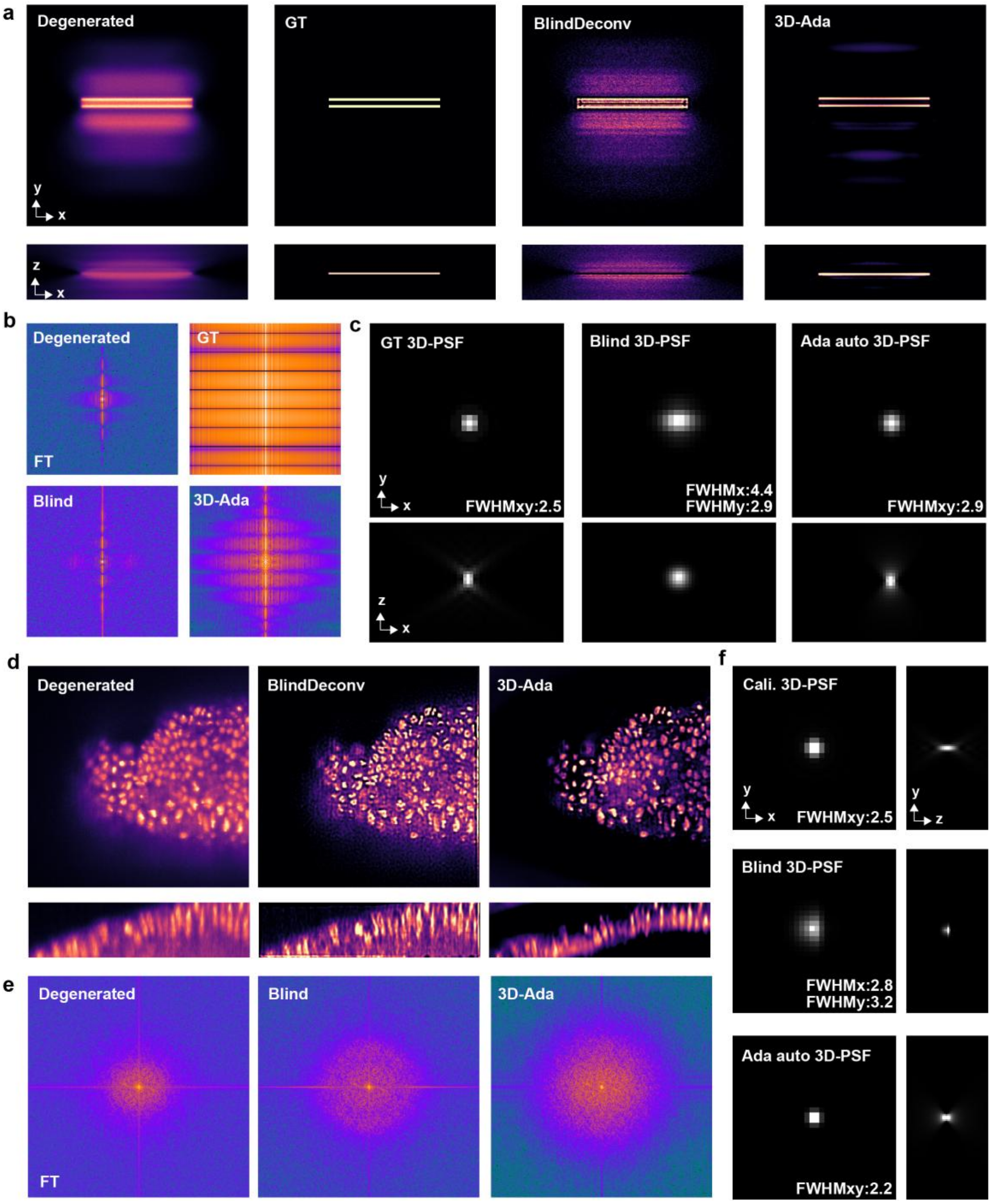


**Fig. S5. Examination of the automatic 3D-PSF calibration methods using the EPFL open-source dataset.** a. A bar pattern and the blind 3D-RL and 3D-Ada deconvolution results. b. The Fourier spectra. c. The 3D-PSF estimated by our scheme, blind 3D-RL deconvolution, and the ground-truth 3D-PSF. d. A Drosophila image and the blind 3D-RL and 3D-Ada deconvolution results. e. The Fourier spectra. f. The 3D-PSF estimated by our scheme, blind 3D-RL deconvolution, and the calibrated 3D-PSF.

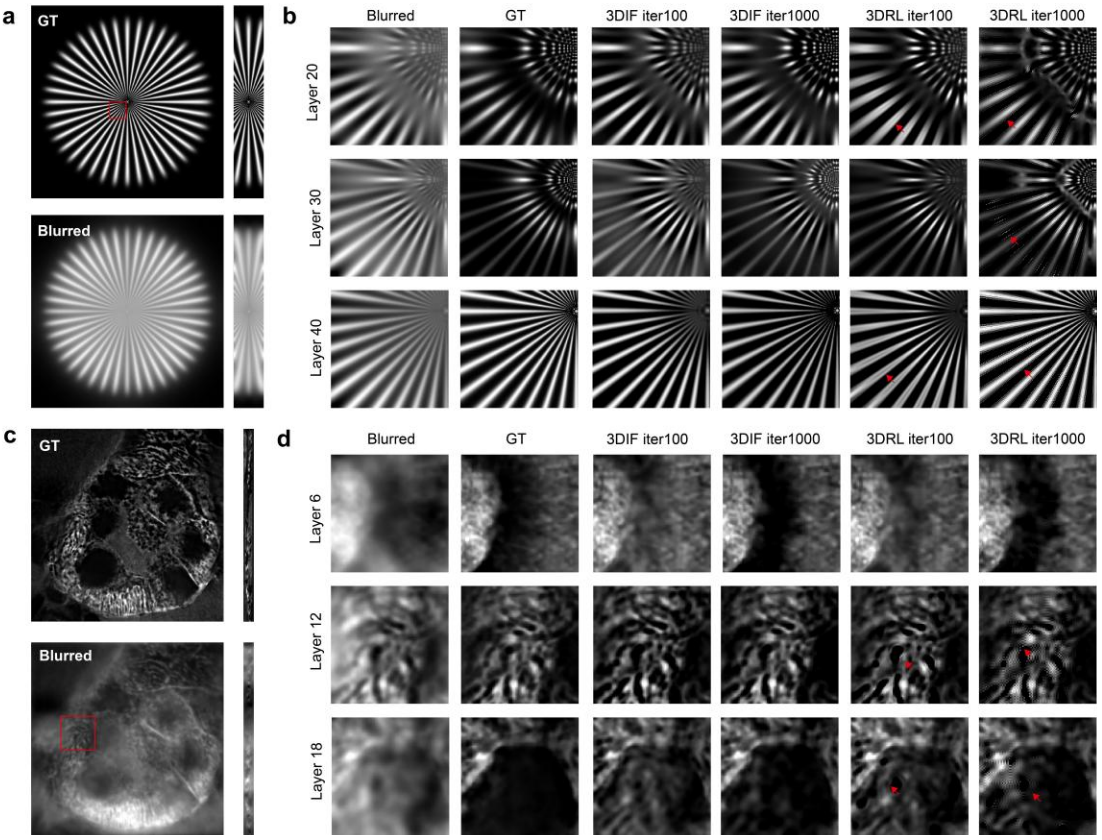


**Fig. S6. Comparison of energy-based 3D inverse-filtering and statistics-based 3D RL in eliminating the blurring caused by 3D PSF. a.** The simulated volumetric data and its blurred version. **b.** Deblurring results of a. using different strategies. **c.** The captured volumetric data of the mouse kidney section and its blurred version. **d.** Deblurring results of c. using different strategies.

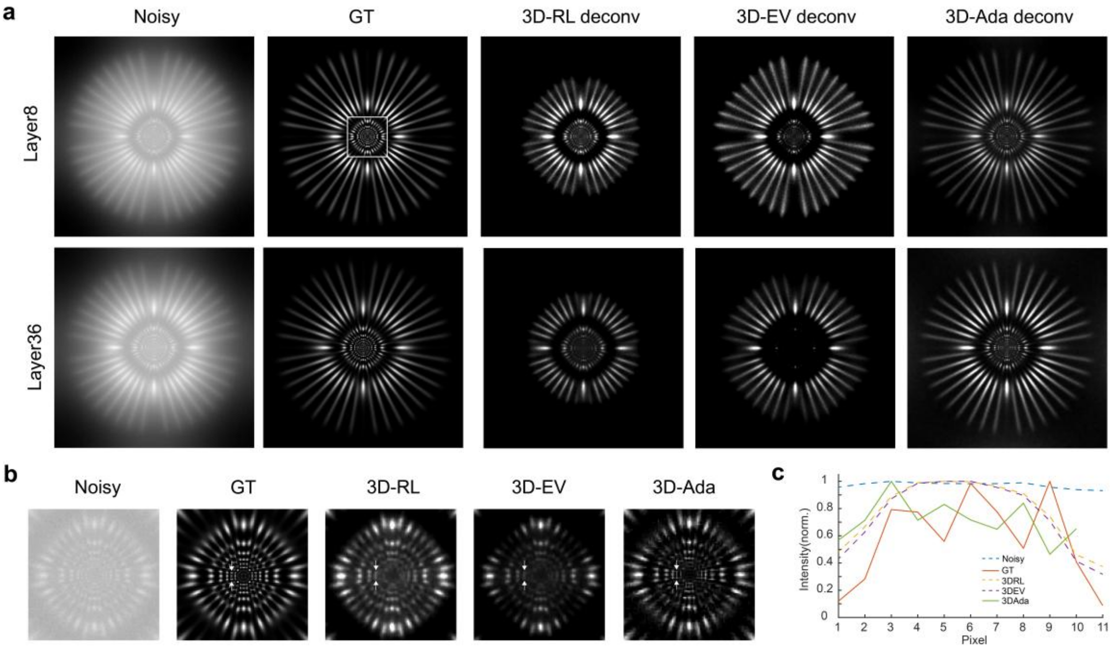


**Fig. S7. Comparisons of 3D-Ada deconvolution with 3D-RL and 3D-EV deconvolution. a.** Two representative layers of a simulated volumetric pattern. **b.** Magnified white boxed region in a. **c.** The intensity profiles along the dotted line in b.

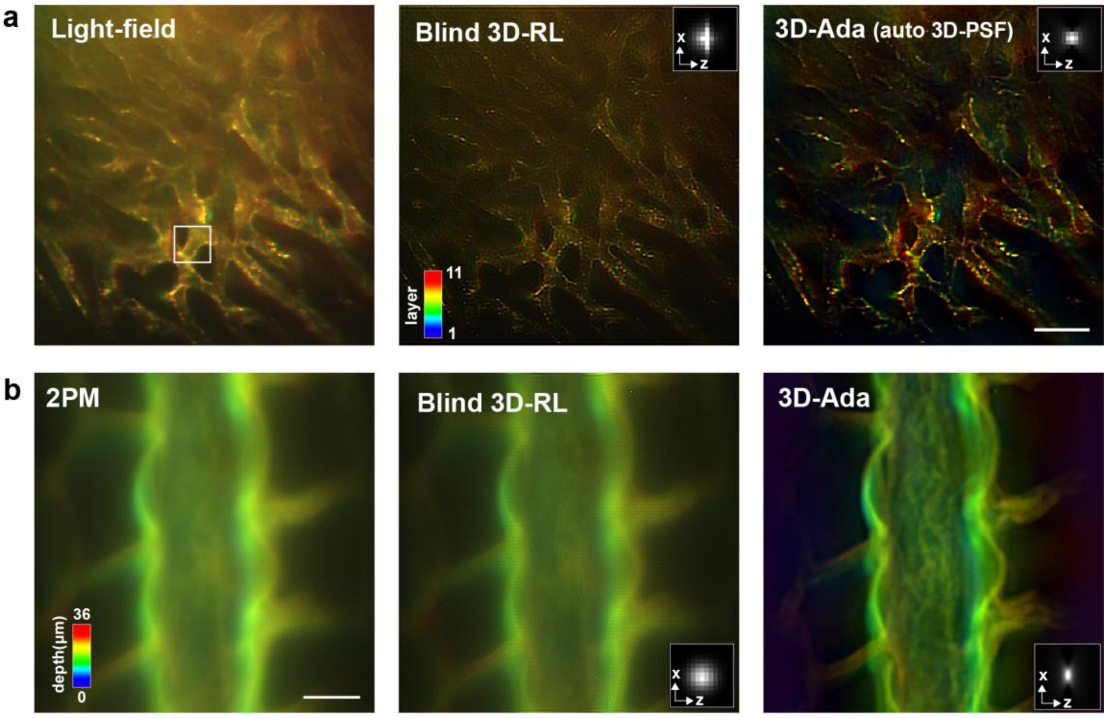


**Fig. S8. Comparison of the 3D-Ada deconvolution and the blind deconvolution on recovering the a.** light-field stack and **b.** two-photon microscope stack. Scale bars: 100 $\mu$m (a), 10 $\mu$m (b).

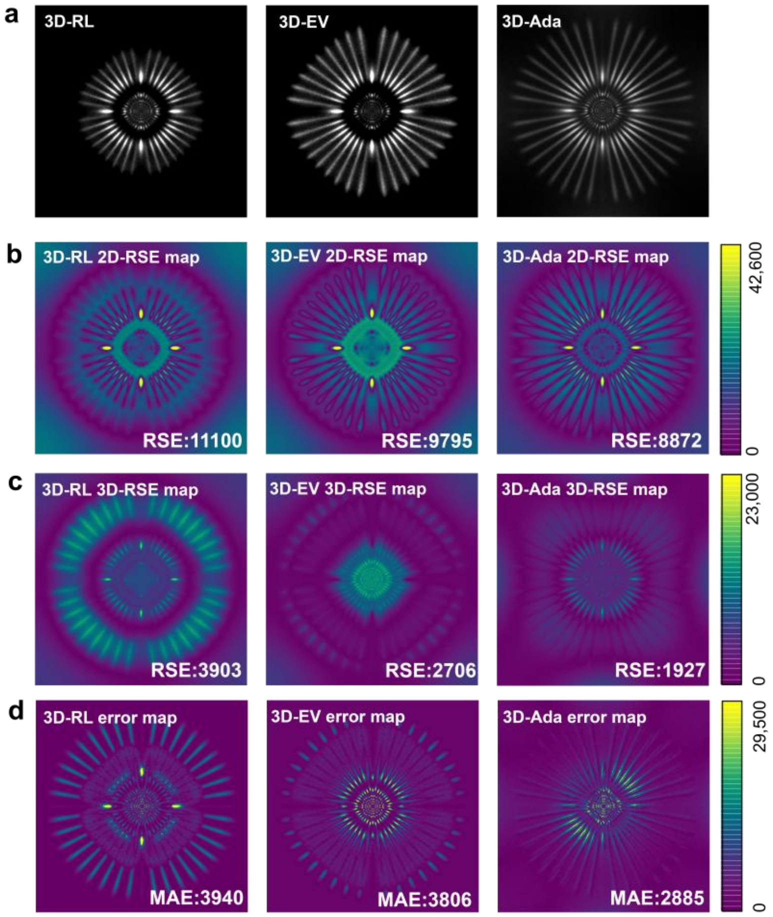


**Fig. S9. 3D-SQUIRREL helps assess 3D deconvolution results. a**. The deconvolution result of layer 8 in Fig. S3. **b.** The 2D-RSE maps of different deconvolution results obtained via the NanoJ-SQUIREL algorithm. **c.** The 3D-RSE maps were calculated using the method proposed in this study. **d.** The GT-error maps. The results show that 2D-SQUIREL would wrongly assess axial resolution enhancement as an error. While 3D-SQUIREL can give an objective assessment that is close to ground-truth error.

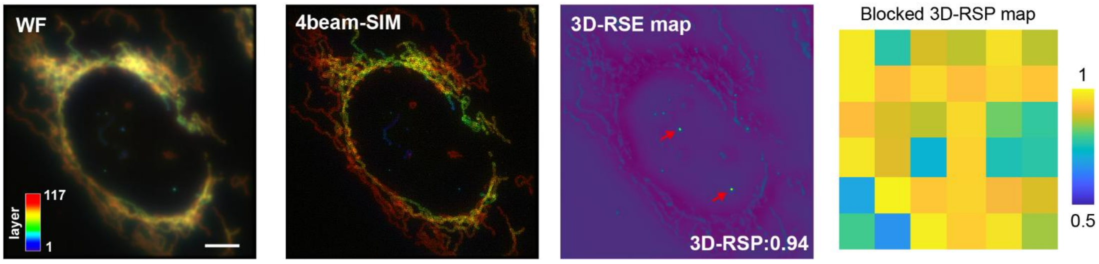


**Fig. S10. 3D-SQUIRREL helps assess physical 3D super-resolution quality.** Assessing the 3D super-resolution quality in the 4beam-SIM system, using the provided open-source images of mitochondria. Scale bar: 10 $\mu$m.

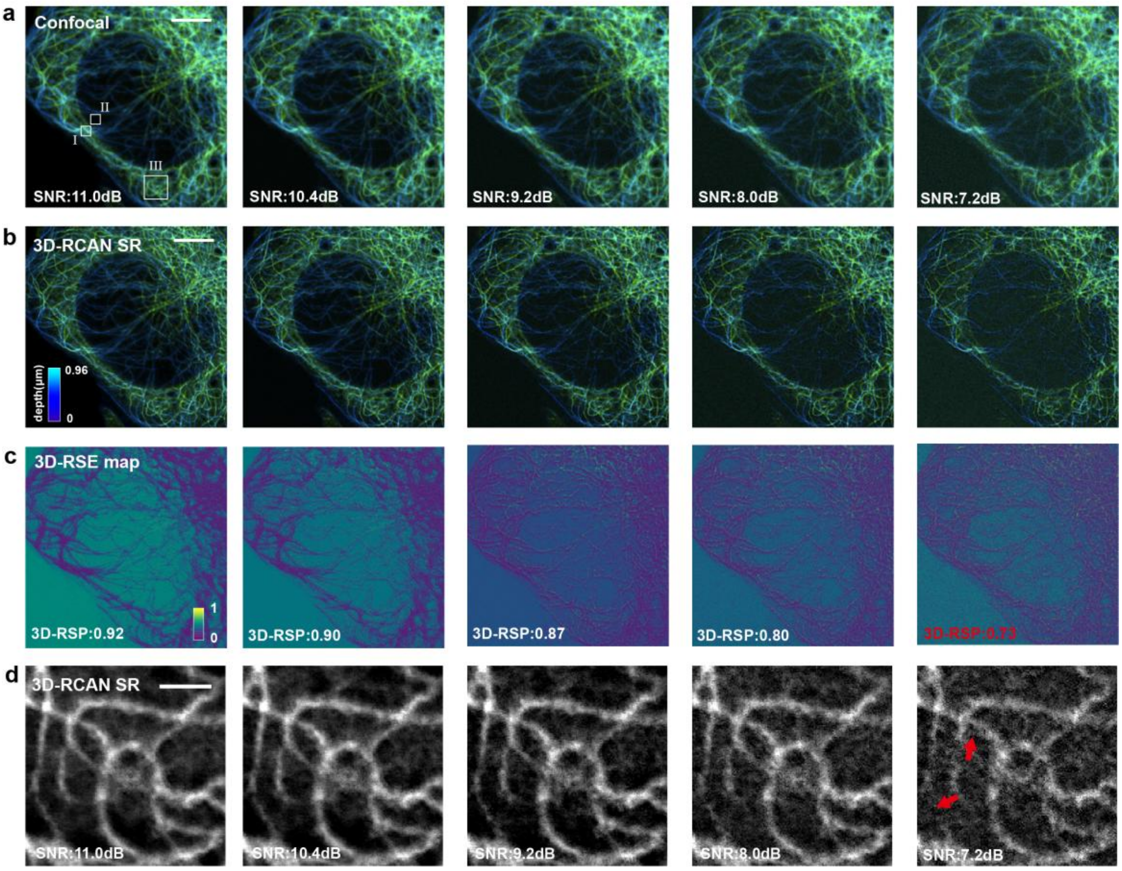


**Fig. S11. Assessing the reliability of deep-learning-based 3D-SR results without knowing the ground-truth. a.** Raw confocal images with different SNR levels. Regions I and II are used to estimate signal and noise intensity, respectively. **b.** The 3D-RCAN SR results of confocal images with different SNR levels. **c.** The 3D-RSE maps of each 3D-SR result obtained with our 3D-SQUIRREL algorithm. **d.** Magnified region III in a. Scale bars: 5 $\mu$m (a, b), 1 $\mu$m (d).

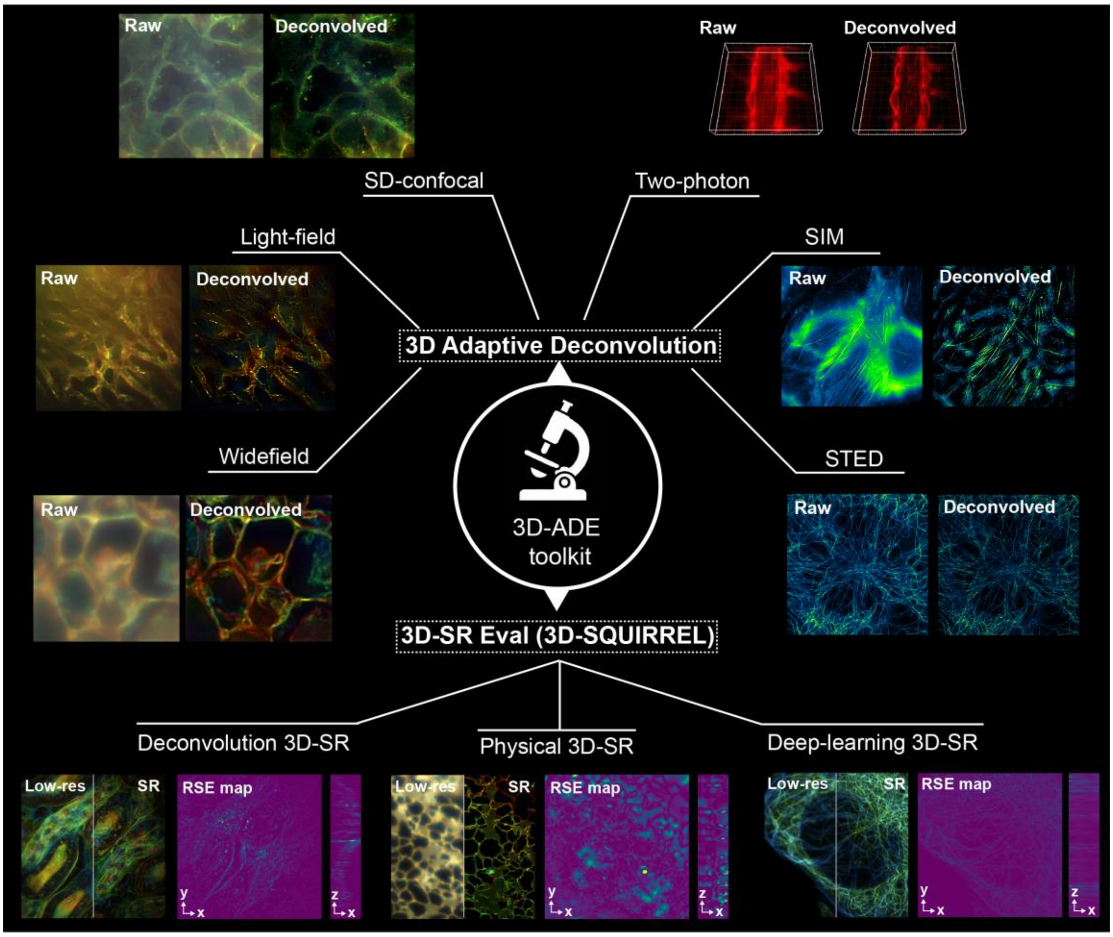


**Fig. S12. A summary of the 3D-ADE toolkit for 3D fluorescence microscopy.**
